# Model-driven generation of artificial yeast promoters

**DOI:** 10.1101/748616

**Authors:** Benjamin J. Kotopka, Christina D. Smolke

## Abstract

Promoters play a central role in controlling gene regulation; however, a small set of promoters is used for most genetic construct design in the yeast *Saccharomyces cerevisiae*. Generating and utilizing models that accurately predict protein expression from promoter sequences would enable rapid generation of novel useful promoters and facilitate synthetic biology efforts in this model organism. We measured the gene expression activity of over 675,000 unique sequences in a constitutive promoter library, and over 327,000 sequences in an inducible promoter library. Training an ensemble of convolutional neural networks jointly on the two datasets enabled very high (R^2^ > 0.79) predictive accuracies on multiple sequence-activity prediction tasks. We developed model-guided design strategies which yielded large, sequence-diverse sets of novel promoters exhibiting activities similar to current best-in-class sequences. In addition to providing large sets of new promoters, our results show the value of model-guided design as an approach for generating useful DNA parts.

## 1. Introduction

Promoters play a central role in the regulation of protein expression, a key task for both natural and engineered biological systems. In the context of bioengineered systems, precise control of gene expression is critical for tasks such as balancing enzyme expression levels in engineered metabolic pathways^1–3^ and building gene circuits to control cell behavior based on external stimuli^4, 5^. Thus, the availability of large sets of promoters with useful properties has the potential to advance the design of sophisticated genetic constructs.

Synthetic biology applications have largely focused on a small number of model systems. The model yeast *Saccharomyces cerevisiae* is well-characterized and relatively straightforward to genetically engineer^6^; as such, it is frequently studied in applications spanning microbial biomanufacturing^7, 8^, synthetic genomes^9^, and genetic circuit design^5, 10^. At present, genetic construct design in this organism generally relies on a small number of well-characterized, native promoters. While additional useful sequences have been uncovered by “mining” the yeast genome,^11^ natural genomes contain a limited number of strong promoters that lead to high levels of protein production in diverse environments. Methods for constructing artificial promoters with desired properties offer a compelling alternative to the existing approach of harvesting natural promoter elements. Typically, sequence-diverse^2^ and short^12^ sequences with high transcriptional activity are desired. Previous efforts have generated artificial promoter libraries through mutagenesis of a wild-type template^13^ or through assembly and screening of random sequence libraries^12, 14^. As another approach, natural promoters can be rationally engineered to modify their properties; for example, binding sites for the artificial transcription factor ZEV, which induces gene expression in the presence of beta-estradiol, were introduced into the yeast P_GAL1_ and P_CYC1_ promoters to create a set of orthogonally inducible sequences^15^. However, strategies based on diversifying sequences from native promoters can encounter drawbacks. Mutagenesis-derived promoters have similar sequences, which may lead to challenges in assembly when using homology-guided sequence assembly methods^16^, and assembly from random sequences can require screening tens of millions of constructs to identify a handful of promoter sequences with desired properties^12^.

In contrast, model-guided approaches to sequence design offer the promise of delivering made-to-order sequences exhibiting specified biological properties. In one example, a Hidden Markov Model predicting nucleosome occupancy from sequence^17^ based on a map of nucleosome positions in the native yeast genome^18^ was used to redesign native yeast promoters and artificial promoters for higher expression^19^. However, such approaches generally require datasets in which the function of interest is measured for a large set of sequences to train a model capable of predicting function from DNA sequence.

When modeling promoter activity, the required data is best acquired by constructing and characterizing libraries of artificial constructs. Massively parallel reporter assays (MPRAs), which characterize the activity of large (usually 10^5^-10^8^) libraries of DNA sequences, can provide the large datasets required to train complex models. FACS-seq is a well-established MPRA^20, 21^ for measuring gene-regulatory activities across entire libraries in a single fluorescence-activated cell sorting (FACS) sort and next-generation sequencing experiment^20, 21^. In this technique, a library is sorted by the abundance of a fluorescent protein whose production is controlled by the promoter (or regulatory element) of interest. The distribution of cells detected in each sort bin is then used to generate a quantitative estimate of protein production. FACS-seq has been used to characterize libraries of randomized 5’ UTRs^22^ and short, complete artificial promoters^23, 24^ in yeast. These large datasets were used to investigate the effect on promoter activity of hand-selected sequence properties^22–24^.

Additionally, researchers have recently attempted to build models capable of predicting promoter activity directly from sequence, without predetermining the features of interest. Deep learning techniques such as convolutional neural networks (CNNs) have been shown to perform well on modeling tasks using large-scale genomics data^25, 26^. CNNs were applied to modeling yeast MPRA datasets^27^, and CNNs trained on MPRA datasets of artificial 5’ UTRs were exploited to design novel functional 5’ UTRs in both yeast and human cells^27, 28^. However, fully exploiting this modeling approach for synthetic biology applications requires extending it to full-scale promoter design, which requires modeling a longer and more complex sequence, making data collection and modeling more challenging.

To model and design entire novel promoters, we adopted a strategy which borrows conserved motifs from known promoters and seeks to learn a sequence-function relationship for the intervening spacer sequences between these motifs. We performed FACS-seq on two libraries comprising over 675,000 full-length constitutive promoters and over 327,000 ZEV-inducible promoters. Using these large datasets, we developed predictive models of promoter activity with an accuracy comparable to state-of-the art models of only the 5’ UTR. We then implemented sequence design strategies using the models’ predictions to generate large, sequence-diverse promoter sets, which we confirmed to be highly active *in vivo*. *In silico* mutagenesis of designed sequences elucidated sequence features identified as significant by the model. Our work provides a set of new promoters with useful properties for synthetic biology applications, as well as a tool for generating promoters with user–specified functional properties. More broadly, our work demonstrates the value of CNNs trained on MPRA-generated data as a tool for designing complex biological sequences with user-specified properties.

## 2. Results

### 2.1 High-throughput characterization of complex promoter libraries

We sought to develop a model capable of accurately predicting promoter activity from sequence in order to generate promoters with specified activities. To generate a dataset for model training, we began by building and characterizing a library of artificial promoters based on the native GPD promoter (P_GPD_; also known as P_TDH3_). P_GPD_ exhibits high activity under standard culture conditions^29^, making it a common choice in the design of genetic constructs and in efforts to engineer synthetic promoters^30^.

Yeast promoters have a modular architecture, which we leveraged in designing the promoter libraries. Functionally important motifs, such as transcription factor binding sites (TFBSes) and the TATA box or other motifs recruiting general transcription factors, are strongly conserved^31^, and while the transcription start site (TSS) can vary^32^, there is evidence that certain motifs are preferred^33, 34^. The sequences between these motifs are not as strongly conserved, but their content still influences promoter activity, particularly the core promoter region extending from the TATA or TATA-like motif to the translation start site^35^. This region includes the 5’ UTR, which has an additional regulatory role at the translational level^36^. Thus, we created a sequence-diverse library by fixing the key conserved motifs in P_GPD_ and varying the surrounding “spacer sequences”. We identified the conserved motifs, including TFBSes for the transcription factors Rap1p and Gcr1p, TATA box, and TSS, through published literature^37^ and the JASPAR transcription factor motif database^38^. Sequence regions outside these motifs were annotated as spacer sequences, and we defined the P_GPD_ promoter as starting approximately 100 bp upstream of the first annotated Rap1p site. The annotated sequence was used as a basis for designing promoter libraries (Supplementary Fig. 1).

Promoter libraries were generated by keeping the identified conserved motifs intact and randomizing the intervening spacer sequences. Where possible, the spacer regions were designed to be shorter than the corresponding sequences in the original GPD promoter to ensure the libraries could be sequenced on a short-read sequencing platform and to increase utility of the final promoter designs. In general, spacer regions were designed as random sequences with the four DNA bases appearing at equal frequencies. However, because the abundance of T nucleotides between the TATA box and TSS is an important predictor of promoter activity^35^, we designed this region such that the bases appear at the same frequencies as in the original P_GPD_ core promoter. Additionally, G was excluded after the TSS motif to prevent out-of-frame premature ATG start codons from being generated which result in weakened activity^22^.

We designed seven P_GPD_ libraries to examine the effect of varying four design parameters (Supplementary Fig. 2): length of the Rap1p binding sites (20 bp for the first site and 18 bp for the second in designs 1, 2, 5, 6; 10 bp each in design 3; absent in designs 4, 7), frequency of each base in the spacer regions between the TFBSes (wild-type base frequency in designs 1, 5, 6; fully randomized in others), length of the spacer between the TFBSes and the TATA box (74 bp in designs 5, 7; 37 bp in others), and length of the TATA motif (10-bp core motif used in designs 6, 7; 18-bp motif in others). The libraries were assembled using a Golden Gate assembly approach and cloned into a two-color plasmid, in which GFP expression was driven by the promoters, and mCherry expression was driven by a constitutive control promoter, P_TEF1_(Fig. 1A). To control for noise in gene expression, the ratio of GFP to mCherry expression levels in each cell was taken as a measure of gene expression activity for a promoter^39^ (Supplementary Fig. 3).

**Figure 1:**
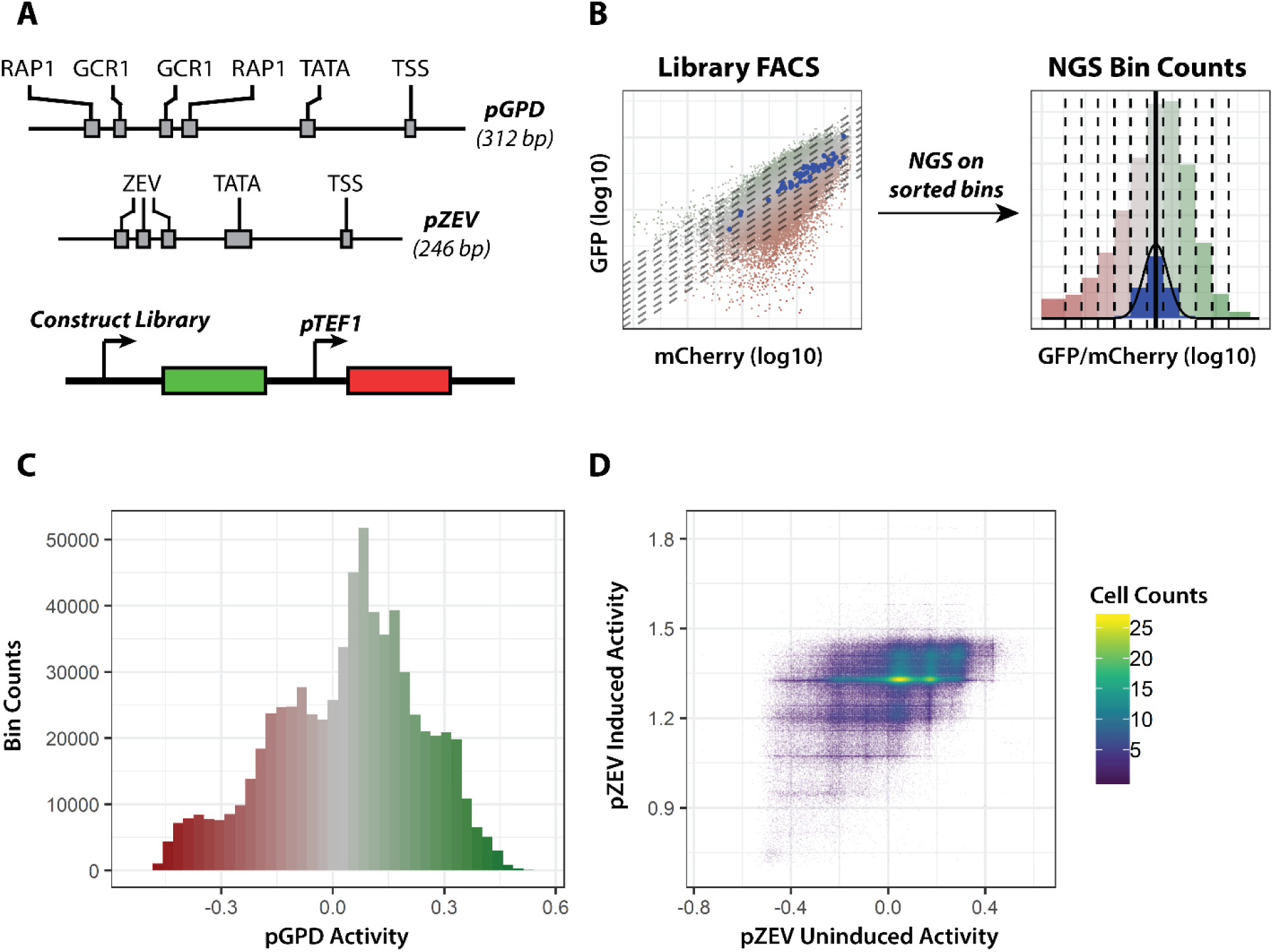
FACS-seq experimental strategy and dataset overview. **A:** Schematic of tested libraries (*above*), indicating regions held constant in promoter design (*grey boxes*); schematic of two-color reporter device used to characterize promoter activity (*below*). “RAP1”, “GCR1”, “ZEV”: transcription factor binding sites; “TATA”: TATA box motif; “TSS”: transcription start site motif. **B:** Schematic of FACS-seq approach for high-throughput promoter activity characterization, in which next-generation sequencing (NGS)-derived histograms of sequence counts in FACS bins generated by sorting a library on promoter activity are used to derive promoter activity for each sequence in a library. **C:** Histogram of promoter activities (log10 ratio of mean GFP to mCherry intensity, in arbitrary units) in the final P_GPD_ library. Only sequences for which at least 10 NextSeq reads were counted in each replicate were used in this analysis. **D:** Density scatter plot of induced and uninduced promoter activities measured in the final P_ZEV_ library. Only sequences for which at least 20 NextSeq reads were counted in each replicate were used in this analysis.

We determined the distribution of gene expression activities from the promoter libraries by transforming library constructs into the strain CSY3 (W303 MATα) assaying each cell population via flow cytometry (Supplementary Fig. 2). With the exception of deleting the Rap1p binding sites entirely, none of the modifications tested had a marked impact on the distribution of library promoter activities. We designed a final library (“Final” in Supplementary Fig. 2) to maximize library diversity (by excluding unnecessary constant regions and using fully randomized spacer sequences when possible) without causing a marked decrease in observed promoter activities. This library was constructed to include 10-bp Rap1p sites and selecting short constant regions and long, fully randomized spacers from the tested design choices. The final promoter library exhibited gene expression activities that span over an order of magnitude, with the highest activity sequences exhibiting levels similar to P_TEF1_, or approximately three times lower than P_GPD_ (Supplementary Fig. 2). The final promoter library encoded promoters that were 312 bp in length, with 83% of the sequence being randomized (Supplementary Fig. 4).

We took a similar approach to the design of inducible promoter libraries. We designed four libraries based on the ZEV promoter system^40^, which relies on an artificial transcription factor (ATF) to induce expression from promoter sequences containing the ATF binding motif in the presence of beta-estradiol. We designed, built, and characterized four P_ZEV_ libraries, varying the number of ZEV ATF binding sites (5 sites in designs 1, 2; 3 sites in others), length of the internal spacer (37 bp in designs 1, 4; 74 bp in others), and length of the TATA box (18 bp in designs 1, 4; 10 bp in others) (Supplementary Fig. 5). We generated a yeast strain expressing the ZEV ATF from the ACT1 promoter (CSY1252), transformed this strain with the ZEV promoter libraries cloned into the two-color characterization plasmid, and characterized promoter activity in the presence of 0, 0.01, and 1 µM beta-estradiol. We observed similar promoter activities in the uninduced state for all designs. Activities in 1 µM beta-estradiol were higher in both designs with the 18-bp TATA box, independent of the number of ZEV sites. We proceeded with design 4 (3 ZEV sites, 37-bp internal spacer, 18-bp TATA box) as the final promoter design to incorporate fewer conserved regions. The final promoter library encoded promoters that are 246-bp in length, with 79% sequence randomized (Supplementary Fig. 6).

We used FACS-seq^20^ to measure the activities of individual promoter sequences in the final P_GPD_ and P_ZEV_ promoter libraries. Each library was sorted into 12 bins based on the ratio of GFP to mCherry fluorescence (Fig. 1B). Preliminary experiments and prior studies^20^ indicated that the GFP:mCherry ratio follows a log-normal distribution with a uniform variance for individual sequences within a library. We refer to the base 10 logarithm of this ratio as promoter activity. Promoters from each sorting bin were recovered by plasmid extraction, tagged with a unique identifier barcode by bin, and analyzed through NGS. The distribution of counts across the 12 bins was used to estimate promoter activity for each library sequence (see Methods). The activities of over 700,000 promoter sequences in the P_GPD_ library (Fig. 1C) and 328,000 sequences in the P_ZEV_ library (Fig. 1D) were measured using this approach.

The FACS-seq experiment on the final GPD promoter library was performed in duplicate. Sequences for which the replicate promoter activity estimates differed by more than 0.2 or which were only observed in the highest or lowest bins were rejected as outliers. After discarding outliers, final measurements on approximately 675,000 sequences were obtained. Data from the replicate experiments were consistent, with a coefficient of determination (R^2^) of 0.94. The mean of activities calculated in each replicate was used as the final estimate of activity for each sequence. The final P_GPD_ library dataset contains sequences spanning over an order of magnitude of promoter activities (from -0.521 to 0.560, median of 0.064, interquartile range of 0.273) (Fig. 1C, Supplementary Fig. 7).

The FACS-seq experiment on the final ZEV promoter library was performed in the presence and absence of 1 µM beta-estradiol. Sequences for which all observed reads fell in either the highest or lowest bins in either the uninduced or the induced condition were discarded as outliers, leaving approximately 327,000 sequences in the final P_ZEV_ dataset (Fig. 1D). In the uninduced case, measured activities ranged from to -0.76 to 0.62 (median of 0.04, interquartile range of 0.272). In the induced case, measured activities ranged from 0.69 to 1.84 (median of 1.33, interquartile range of 0.122).

### 2.2 A convolutional neural network accurately predicts promoter activity

We next sought to leverage the promoter sequence-activity datasets to build a model that predicts promoter activity. Rather than attempting to hand-select the sequence features responsible for determining promoter activity, we modeled activity as a function of raw sequence by implementing a convolutional neural network (CNN) which accepts a one-hot encoded DNA sequence as input and outputs a quantitative activity prediction. The model consists of a series of convolutional and max-pooling layers, which learn a compressed representation of salient features within the input sequences. The output of these layers is fed into a two-layer fully-connected network, which outputs a single numerical prediction of promoter activity. All convolutional layers had a width of 8 and 128 output channels and used a rectified linear unit (ReLU) nonlinearity, with batch normalization applied between the convolution and the ReLU layer. Input sequences were one-hot encoded and processed with six rounds of convolution and max-pooling with a stride length of 2. Two 128-unit fully connected layers, each followed by batch normalization and a ReLU, were then applied, followed by a final fully-connected layer which directly provided the final output(s). L2 regularization with a weight of 10^−4^ was applied at all layers except the final output.

To model promoter activity in the P_GPD_ library, the P_GPD_ library data was divided into training (80% of sequences), validation (10%), and test sets (10%). The model was trained on the training dataset, with performance on the validation data monitored to avoid overfitting. Training was interrupted after five epochs passed without improved performance on the validation data. A final model was obtained after 18 training epochs (Fig. 2A) and used to generate promoter activity predictions on the test dataset. The predictions accounted for 65% of the variability in the test data (Fig. 2B), suggesting that the model’s predictions were generalizable to sequences not included in the training data.

**Figure 2:**
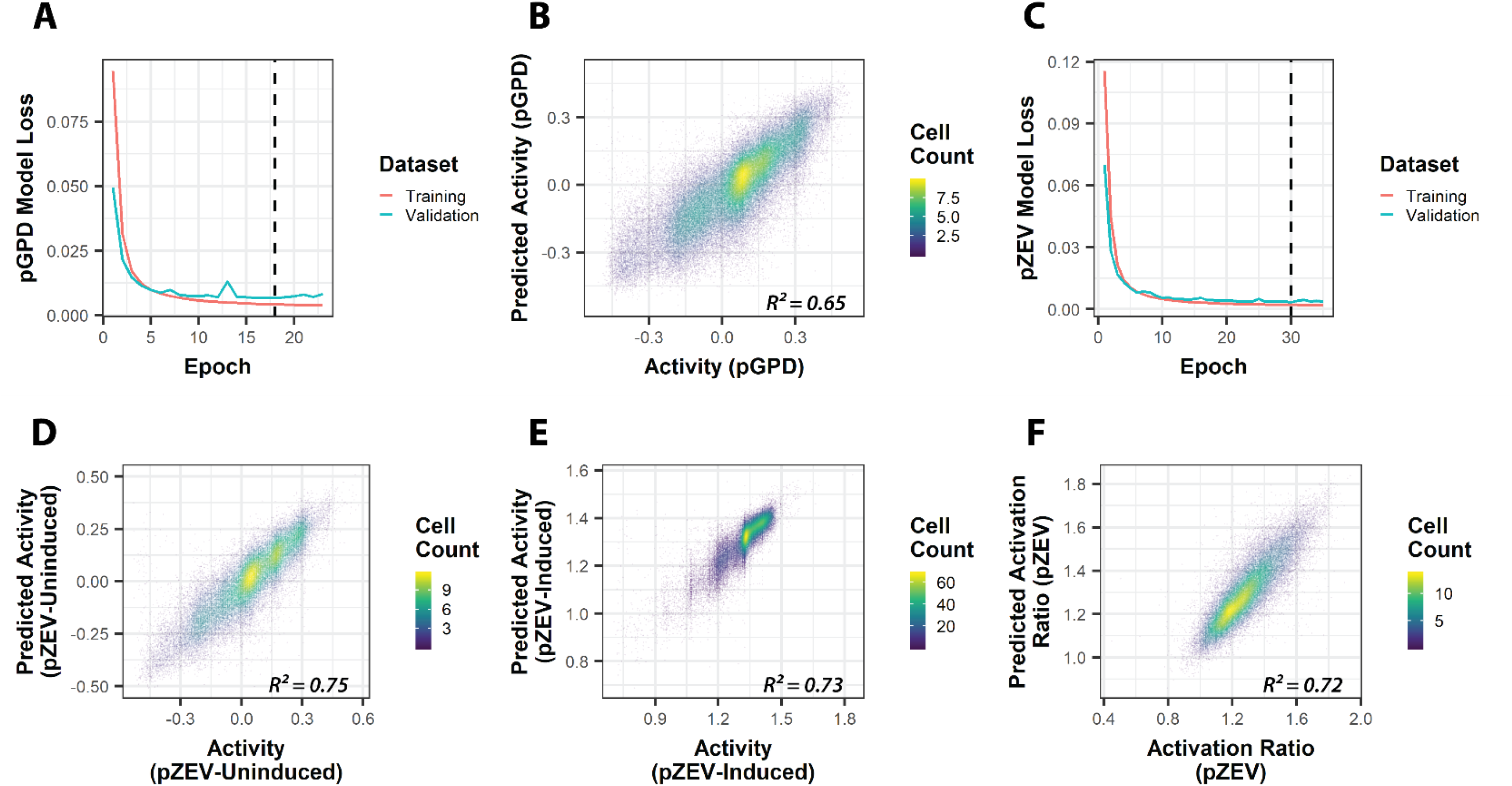
Neural networks trained on P_GPD_ and P_ZEV_ data accurately predict promoter activity. Only sequences for which at least 10 NextSeq reads were counted in each replicate were used in analyses of P_GPD_ data; only sequences for which at least 20 NextSeq reads were counted in each replicate were used in analyses of P_ZEV_ data. **A:** Model loss curve for P_GPD_ training; dashed line indicates epoch selected by early stopping for the final model. **B:** Predicted promoter activities versus FACS-seq measurements for held-out test data in the P_GPD_ dataset. **C:** Model loss curve for P_ZEV_ training; dashed line indicates epoch selected by early stopping for the final model. **D:** Predicted promoter activities in the uninduced condition versus FACS-seq measurements for held-out test data in the P_ZEV_ dataset. **E:** Predicted promoter activities in the induced condition versus FACS-seq measurements for held-out test data in the P_ZEV_ dataset. **F:** Predicted activation ratios (ratio of predicted induced and uninduced promoter activities) versus FACS-seq-derived activation ratios for held-out test data in the P_ZEV_ dataset.

We next built a model using the same architecture and training process to predict promoter activity in the P_ZEV_ library. In order to generate separate predictions for both the uninduced and the induced conditions in one model, we used an output layer with two units (as opposed to one unit in the P_GPD_ network). The accuracy of both predictions was weighted equally in training. We similarly divided the P_ZEV_ data into training (80%), validation (10%), and testing (10%) datasets, and monitored performance on the validation data to determine when to halt training. A final model was obtained after 30 training epochs (Fig. 2C). We generated predictions from this model on the test dataset and found that P_ZEV_ predictions generalized well to test data, achieving an R^2^ of 0.75 on the uninduced data (Fig. 2D) and 0.73 on induced data (Fig. 2E). Additionally, we calculated a predicted activation ratio from the predicted induced and uninduced activities for held-out P_ZEV_ test data (Fig. 2F). The model performed well, with an R^2^ of 0.72, suggesting that our CNN’s predictions of uninduced and induced activity were robust enough to yield an accurate prediction of activation ratio.

Finally, we examined whether our CNN architecture was sufficiently flexible to jointly model the P_GPD_ and P_ZEV_ datasets. Deep learning is known to perform best on very large datasets, including in genomics applications^41, 42^. As the P_GPD_ and P_ZEV_ promoter designs are nearly identical (apart from the TFBSes and immediately surrounding sequence), we hypothesized that the datasets could be merged and used to train a single model, taking advantage of the increased dataset size. Additionally, we attempted to improve the model’s performance by training an ensemble of models on slightly different datasets; nine sub-models were trained in total. (Loss traces for these models appear in Supplementary Fig. 8.) All sub-models used the same held-out test data, but the remaining data was divided into nine partitions, and each sub-model used a different partition of this data for validation. Final predictions on test data were arrived at by averaging the sub-models’ predictions on the test dataset.

The results show that the “joined” model outperformed the original P_GPD_ and P_ZEV_ models, achieving better fits than the original models on test data for all attempted tasks, including modeling P_GPD_ data (Fig. 3A), P_ZEV_ uninduced (Fig. 3B) or induced (Fig. 3C) data, and the P_ZEV_ activation ratio (Fig. 3D). We computed R^2^ for the sub-models’ individual predictions and for the joined model on each of these quantities. Comparing the R^2^ achieved by the original models to the median R^2^ achieved by the ensemble of sub-models and to that of the final joined model showed that both improvements tested – merging the datasets and averaging the predictions of the model ensemble – contributed to the performance of the final model, which achieved an R^2^ of 0.79 or better on all tested tasks (Table 1).

**Figure 3:**
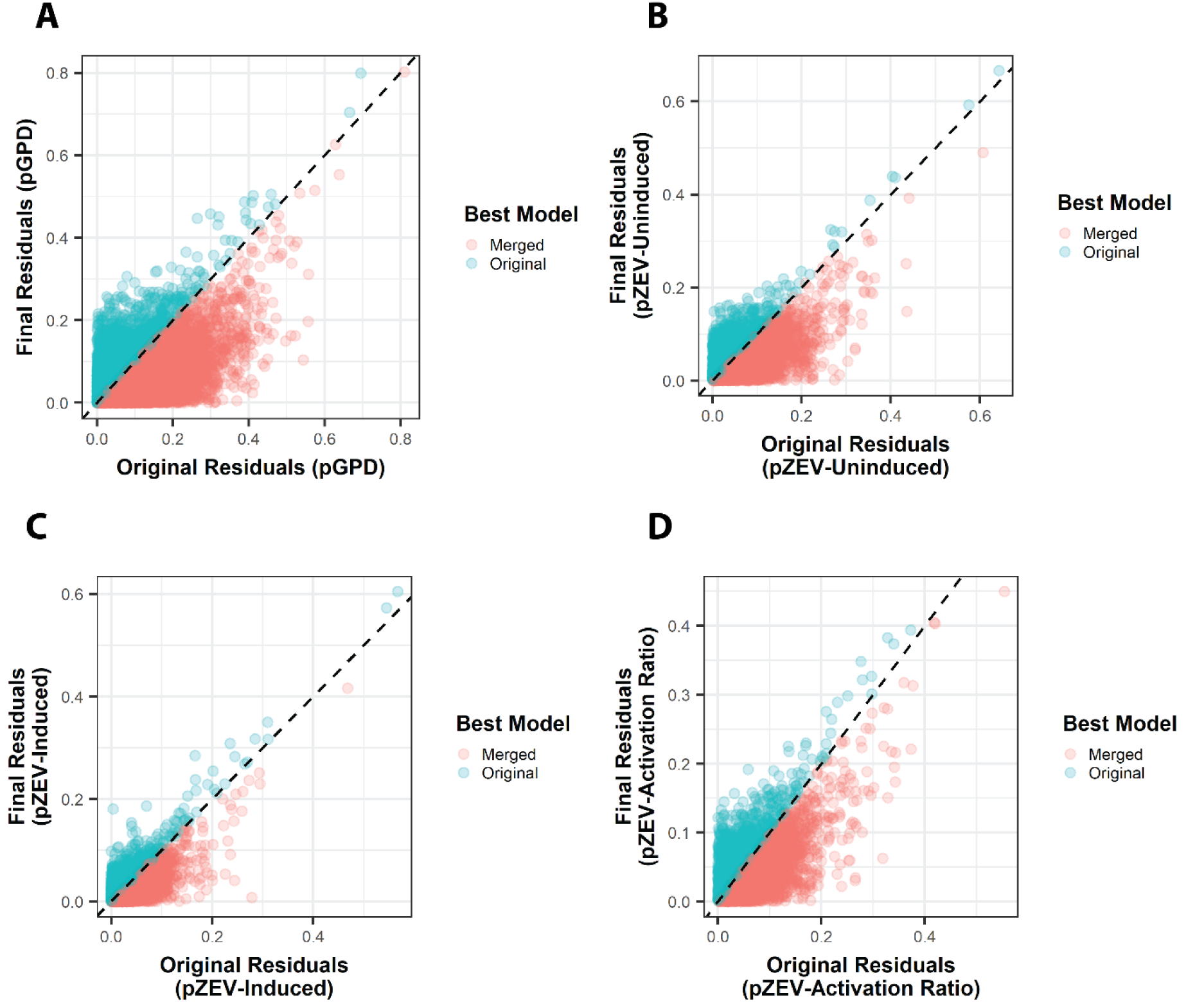
Improving predictions of promoter activity with a model ensemble. In all panels, point color indicates whether the original model (*blue*) or the merged model (*red*) performed better for each measured promoter, as measured by the absolute value of modeling residuals (difference between predicted and measured promoter activities) for the specified data. Dashed line indicates equal residuals between the two models (as a guide for the eye). **A:** Absolute values of modeling residuals for P_GPD_ data, comparing the “final model” – the ensemble of nine submodels, trained jointly on the P_GPD_ and P_ZEV_ datasets – with the original P_GPD_ model (Fig. 2A-B). **B:** Absolute values of modeling residuals for P_ZEV_ data in the uninduced condition, comparing the “final model” with the original P_ZEV_ model (Fig. 2C-F). **C:** Absolute values of modeling residuals for P_ZEV_ data in the induced condition, comparing the “final model” with the original P_ZEV_ model. **D:** Absolute values of modeling residuals for activation ratios (predicted by taking the ratio of predicted induced to predicted uninduced promoter activity) for P_ZEV_ data, comparing the “final model” with the original P_ZEV_ model.

**Table 1:** Coefficients of determination for model fits. The column “Task” describes the data being modeled; “P_GPD_” is as in Figs. 2B, 3A; “P_ZEV_ – Uninduced” is as in Figs. 2D, 3B; “P_ZEV_ – Induced” is as in Figs. 2E, 3C; and “P_ZEV_ – Activation Ratio” is an in Figs. 2F, 3D. Column “Original” contains R^2^ values for the original models (see Fig. 2), column “Median Joined” contains the median R^2^ values for the ensemble of nine submodels when carrying out each prediction task independently, and column “Merged” contains the R^2^ values achieved by taking the average of submodel predictions as a final prediction.

| Task | Original | Median Joined | Merged |
| --- | --- | --- | --- |
| $P_{\text{GPD}}$ | 0.65 | 0.72 | 0.80 |
| $P_{\text{ZEV}} - \text{Uninduced}$ | 0.75 | 0.80 | 0.84 |
| $P_{\text{ZEV}} - \text{Induced}$ | 0.73 | 0.75 | 0.79 |
| $P_{\text{ZEV}} - \text{Activation Ratio}$ | 0.72 | 0.77 | 0.82 |

### 2.3 Model-guided design provides novel promoters with high activities

To validate model predictions and generate new promoters, we developed and tested a set of model-guided sequence design strategies. The models were used to design promoters maximizing one of three objectives: promoter activity (P_GPD_ sequences), activity in the induced condition (P_ZEV_ sequences), and activation ratio (P_ZEV_ sequences). We focused on maximizing these properties through the model, as sequences with lower activities can readily be selected from the original P_GPD_ or P_ZEV_ datasets. Thus, using the model to design sets of constitutive or inducible promoters with high activities will expand the range of promoter properties readily accessible from FACS-seq analysis of the promoter libraries.

Three sequence-design strategies that rely on activity predictions generated by our model were developed: (1) screening, (2) evolution, and (3) gradient ascent. In the screening strategy, random sequences were generated following the specification of the original libraries used to generate the FACS-seq data, and accepted if their predicted values for the objective property, or scores, are above a set threshold (Supplementary Fig. 9). In the evolution strategy, a set of mutagenized variants was generated from a candidate sequence. The variant with the highest predicted activity was accepted if its score is above the threshold. If not, a new set of variants was generated from this sequence, and the evolutionary cycle continues (Supplementary Fig. 10). The gradient ascent strategy is a modification of the iterative gradient descent process used to train neural networks^43, 44^. In the basic implementation of network training by gradient descent, the gradient of all weights in the network with respect to a loss function is calculated, and the weights are updated by subtracting the product of the gradient and a hyperparameter called the learning rate. We implemented a gradient ascent strategy for promoter design by calculating the gradient on the input data itself, with respect to the score property to be maximized. Given a matrix representation of a one-hot encoded DNA sequence, the gradient was iteratively calculated and used to generate an updated version of the input. The objective score was calculated for a “rounded” one-hot matrix derived from this updated input. The sequence was accepted if its score was above a threshold; if not, the gradient ascent process continued (Supplementary Fig. 11).

In addition to testing the three design strategies, we tested the effect of applying an “extrapolation penalty” and/or a “GC constraint” when calculating a final promoter activity estimate from the model ensemble. Merging the predictions of the ensemble’s sub-models by taking their mean carries the risk that an outlier misprediction from a single sub-model results in an inaccurate estimate of promoter activity. To generate conservative predictions by imposing an extrapolation penalty, we merged the sub-models’ predictions by computing the mean of predictions minus their standard deviation. The GC constraint imposes the condition that no 20-bp window in a sequence has a GC content of less than 25% or greater than 80%. As extremes in GC content can make common molecular biology procedures like PCR and Sanger sequencing challenging^45^, the GC constraint ensures resulting promoter designs are tractable for downstream applications. This approach also ensures that our novel promoters differed from the wild-type P_GPD_ promoter, as the 141-bp core promoter sequence has a GC content of approximately 23%, including a 60-bp stretch with a GC content below 17%.

We designed a total of 33 design approaches applying various combinations of these strategies and objectives, as well as the extrapolation penalty and the GC constraint (Supplementary Table 1). For each of our three objectives (P_GPD_-Activity, P_ZEV_-Induced activity, P_ZEV_-Activation Ratio), the design threshold for the evolution and gradient ascent strategies was set such that a promoter set of 120 sequences could be generated within an hour for a reference design using the evolution strategy, merging the model outputs using the mean, and not applying the GC constraint. Additionally, a design was generated for each objective using the gradient ascent strategy and the extrapolation penalty, not applying the GC constraint, and increasing the threshold in increments of 0.05 until a set of 120 sequences could no longer be generated within an hour. The screening strategy was unable to generate promoter designs that reached the starting thresholds used for evolution and gradient ascent designs. Accordingly, the threshold was decreased in increments of 0.05 until a promoter set could be generated. The GC constraint was not imposed on any of the approaches using the screening strategy. Finally, sequences that contained BsaI restriction sites which interfere with our assembly strategy (see Methods) were removed; in each of the designed promoter sets, more than 100 designs remained.

We characterized the activities of the designed promoters in a FACS-seq experiment. In addition to the designed promoter sets, generated via a Golden Gate Assembly approach, sequences from the original GPD and ZEV promoter libraries were selected for synthesis as controls to validate the dataset quality, examine whether outliers in model predictions resulted from errors in FACS-seq data, and directly compare the best-performing promoter sequences in the original libraries to the model-designed promoters. The criteria for selecting control sequences and the number of sequences in each control promoter set are summarized in Supplementary Table 2. We transformed the resulting library of control and designed promoter sets into CSY1252 and characterized the individual promoter activities using FACS-seq (Supplementary Fig. 12).

We determined how well-represented the designed or control promoter sets were in the experimental data. We set a threshold of at least 20 sequences measured in the FACS-seq experiment for a sequence to be measured. All sets met this threshold except for two of the P_GPD_ control sets (sets 2 and 4, respectively representing a range of P_GPD_ activities and sequences in the P_GPD_ test data with the greatest prediction errors) and three of the P_GPD_ design sets (sets 22, 24, 28, representing one evolution and two gradient design strategies) (Supplementary Table 1, Supplementary Table 2).

The data on the control promoter sets was used to validate the accuracy of the original FACS-seq data for sequences with extreme activity values (“outliers”) and sequences whose activities were not correctly predicted by the model (“inliers”). Activity measurements in the original and the validation experiments for sequences in the outlier sets were not well-correlated, in contrast to those for sequences in inlier sets (Supplementary Fig. 13). The results suggest that outlier values from the FACS-seq characterization of the original libraries were unreliable, supporting our decision to exclude these sequences from modeling. Sequences from non-outlier control promoter sets were accurately measured (Supplementary Fig. 13, set 11, ‘ZEV-grid’), with the exception that adjustments made to bin edges in order to measure highly active P_GPD_ and P_ZEV_-uninduced sequences reduced the sensitivity of mean measurements for P_ZEV_-uninduced sequences with low activity. To determine whether our model-designed sequences outperformed the best sequences from our initial libraries, control promoter sets that exhibited high activities and were accurately measured in the validation FACS-Seq were selected as benchmarks. Specifically, sets 1 (“Non-“outlier” sequences with highest activity”) and 5 (“GPD test sequences accurately predicted to be highly active”) were used for P_GPD_-Activity comparisons, sets 9 (“Non-“outlier” sequences in Induced with highest activity”) and 16 (“ZEV test sequences accurately predicted to be highly active in Induced condition”) were used for P_ZEV_-Induced comparisons, and sets 10 (“Sequences with high activation ratios in ZEV data”) and 17 (“ZEV test sequences accurately predicted to have high activation ratios”) were used for P_ZEV_-Activation Ratio comparisons. The control promoter sets are referred to as “training data” below in comparison to model-designed promoter sets. Additionally, we fit linear models to the measured activities of sequences in the P_ZEV_ experiments measured in the original P_ZEV_ FACS-seq and validation experiments in order to rescale the P_ZEV_ means to correspond with validation results (Supplementary Fig. 14).

We next evaluated the performance of our promoter design strategies, first examining how well promoters performed on the objective they were designed to maximize. For P_GPD_-Activity designs, in addition to the training data sequences described above, we measured activities for promoter sets for all three design strategies (i.e., screening, evolution, gradient ascent), with a minimum of 20 sequences in each set (Fig. 4A, Supplementary Fig. 15). For all combinations of design strategy and choice of GC constraint, promoter sets for which the extrapolation penalty was applied exhibited significantly higher activities than those where it was not applied (p < 0.01 for all comparisons, one-sided Mann-Whitney test (MWT)).

**Figure 4:**
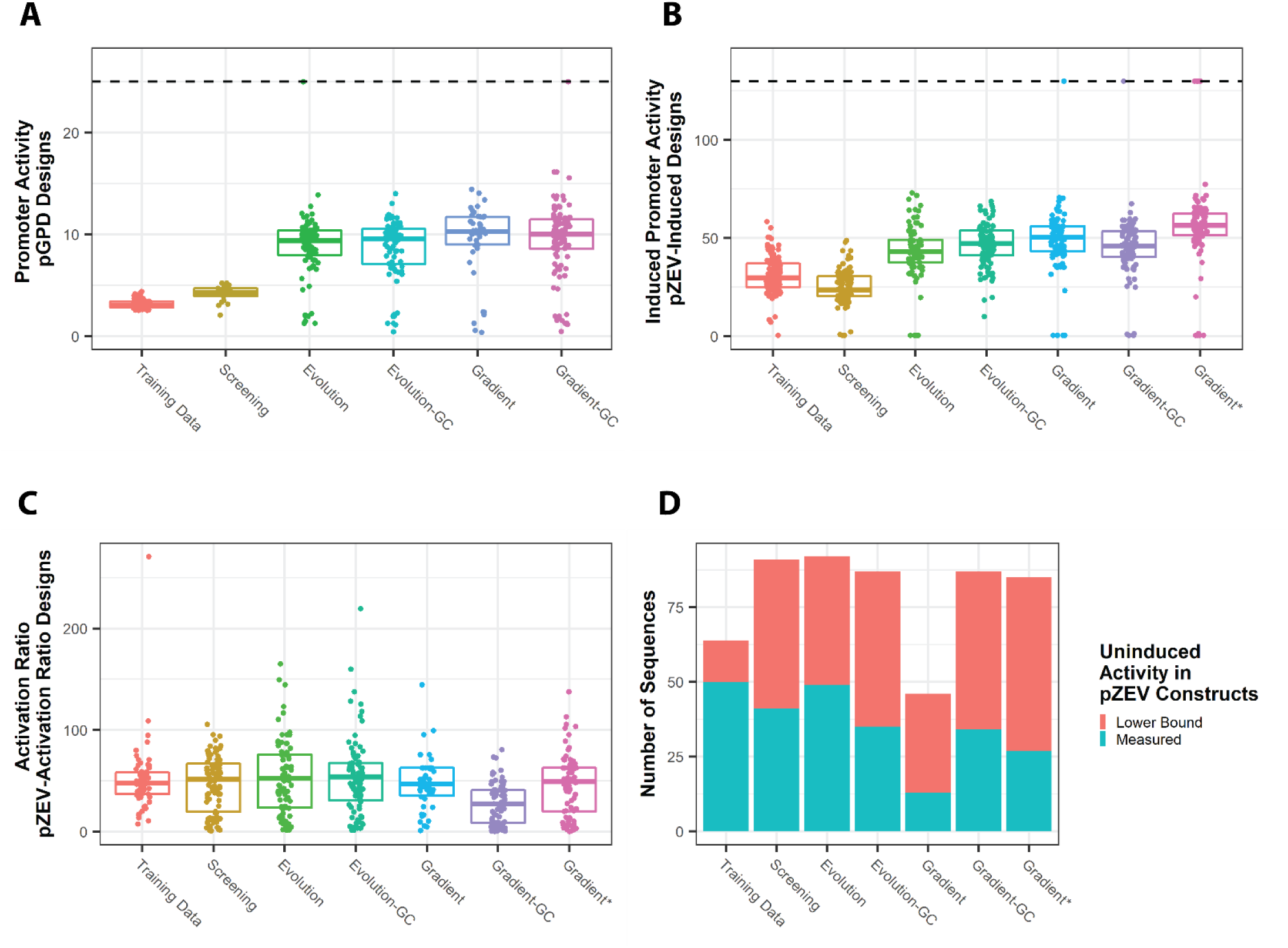
Performance of designed promoter sets in validation FACS-seq experiment. In panels A-C, boxes represent interquartile ranges; the bar within each box indicates the median. Promoter activities are shown here on a linear scale, and were transformed to a scale co-measureable with the results of individual promoter testing using a linear model fit to promoter activities measured by FACS-seq and by individual testing for a set of promoters spanning a range of expression activities. **A:** FACS-seq measurements of promoter activities for P_GPD_ promoter sets (or corresponding training data sequences). “Training Data”: selected highly active sequences from the initial P_GPD_ FACS-seq; “Screening”: P_GPD_ promoter set generated using the screening approach; “Evolution”: P_GPD_ promoter set generated using the evolution approach; “Evolution-GC”: P_GPD_ promoter set generated using the evolution approach, with the GC constraint applied; “Gradient”: P_GPD_ promoter set generated using the gradient ascent approach; “Gradient-GC”: P_GPD_ promoter set generated using the gradient ascent approach, with the GC constraint applied. Points placed along the horizontal line were only measured in the highest-activity bin in FACS-seq. **B:** FACS-seq measurements of promoter activities for P_ZEV_ promoter sets designed to maximize induced activity (or corresponding training data sequences). Axis labels referring to P_ZEV_-Induced sequences and designs, but otherwise as in **A**; “Gradient*”: P_ZEV_-Induced promoter set generated using the gradient approach, with an elevated target threshold set relative to other designs. Points placed along the horizontal line were only measured in the highest-activity bin in FACS-seq. **C:** FACS-seq measurements of promoter activities for P_ZEV_ promoter sets designed to maximize activation ratio (or corresponding training data sequences). Axis labels referring to P_ZEV_-Activation Ratio sequences and designs, but otherwise as in **B**. **D:** Sequence sets displayed in **C**, indicating the number of sequences in each set which were quantitated in the experiment (“Measured”) or which fell entirely in the lowest-activity bin during FACS-seq (“Lower Bound”). All displayed promoter sets were generated using the extrapolation penalty.

We then examined the results across the P_GPD_-Activity promoter design strategies that employ the extrapolation penalty (Fig. 4A). Promoter sequences generated through the screening strategy exhibited higher activities *in vivo* than the training data sequences (median activity, screening: 4.29, training data: 3.03; *p* = 1.15×10^−7^, MWT). Thus, the simplest design strategy generated promoter sequences with higher activities than any sequence present in the training data. In addition, all measured promoter sets generated through the evolution and gradient ascent strategies exhibit substantially higher activities than those generated by the screening strategy. Specifically, the lowest median promoter activity among the evolution and gradient ascent sets (i.e., for evolution designs without GC constraint) was 9.40 versus 4.29 for the screening designs. The gradient ascent strategy generated designs with slightly higher activities than those generated using the evolution strategy (without GC constraint, evolution: 9.40, gradient ascent: 10.27, p = 3.39×10^−3^; with GC constraint, evolution: 9.55, gradient ascent: 10.03, p = 2.16×10^−2^, MWT). Finally, the median sequence activities were almost identical between corresponding promoter sets generated with and without the GC constraint (evolution: 9.55/9.40 with/without GC constraint; gradient ascent: 10.03/10.27 with/without GC constraint).

We next examined whether the model-based design approaches outperformed the training data for ZEV promoters designed to maximize induced activity (Fig. 4B, Supplementary Fig. 16). The extrapolation penalty significantly improved promoter activities for almost all pairs of promoter sets tested with and without the extrapolation penalty (*p* < 0.01 for all comparisons, MWT). One exception to this trend was observed with promoter sets generated using the evolution strategy without the GC constraint, which generated a similar distribution of *in vivo* promoter activities with and without the extrapolation penalty (with/without extrapolation penalty: 43.15/42.76). (Supplementary Fig. 16).

We then examined promoter sets generated for ZEV promoters designed to maximize activity at induced state for all models that use the extrapolation penalty (Fig. 4B). The promoter sequences generated using the screening strategy generally exhibited lower activities than those in the training data (screening: 23.71, training data: 29.80, p = 1.50×10^−6^, MWT). However, all tested evolution and gradient ascent strategies generated promoter sets that exhibit higher activities than those in the training data. In particular, the lowest median promoter activity exhibited across the evolution and gradient ascent designs (for evolution designs without GC constraint) was 43.16 versus 29.80 for training data sequences. For promoter sequences generated without the GC constraint, the gradient ascent approach generated sequences with higher activities than those generated with the evolution approach (evolution: 43.16, gradient ascent: 50.41, p =1.44×10^−4^, MWT). However, these trends in promoter activity between the gradient ascent and evolution approaches were reversed when applying the GC constraint (evolution: 47.29, gradient ascent: 46.01). The data indicate that applying the GC constraint did not result in a consistent impact on the model output. Specifically, applying the GC constraint with the evolution strategy resulted in promoter sequences exhibiting higher activities (with/without GC constraint: 47.29/43.16, p =2.24×10^−2^, MWT), whereas the opposite was observed with the gradient ascent strategy (with/without GC constraint: 46.01/50.41 with, p =6.88×10^−3^, MWT).

We also examined the impact of increasing the design threshold for a promoter set generated using the gradient ascent strategy with the extrapolation penalty and without the GC constraint. When increasing the target prediction value from 1.6 to 1.65, we observed an increase in median promoter activity (original threshold: 50.41, elevated threshold: 56.52, p =1.43×10^−6^, MWT). Thus, these results show that increasing the design threshold allowed the generation of promoter sequences with higher activities, which underscores the ability of our model to generate meaningful promoter activity predictions, even for promoter sets that exhibit activities higher than those represented in the training data.

In contrast to the results for the P_GPD_-Activity and P_ZEV_-Induced promoter designs, when using the models to design promoter sequences for the P_ZEV_-Activation Ratio objective, none of the designed promoter sets’ median activation ratio was observed to be significantly greater than that of the training data sequences (MWT, using a significance threshold of 0.05) (Fig. 4C, Supplementary Fig. 17). We determined that many of the designed sequences intended to optimize the activation ratio objective had uninduced activities at the lower limit of detection (i.e., cells containing these sequences were collected in the lowest-activity bin) (Fig. 4D). Specifically, 14 of 64 (21.9%) training data sequences were at the lower limit of detection, versus 46.7% to 71.7% of the designed sequence sets (Fisher’s exact test; p < 10^−2^ for all). This result suggests that our design strategies had selected sequences with low uninduced activities to maximize the activation ratio.

Finally, we analyzed the GC content and inter-sequence diversity of the promoter sequences designed through our model. We analyzed the maximum and minimum GC content for each designed sequence in 20-bp windows. The analysis shows that sequences designed with the GC filter constraint fell within the GC content specification; however, sequences not designed with this constraint virtually never fall within this specification (Supplementary Fig. 18). The results indicate that the applied GC constraint was sufficiently stringent that sequences which satisfied it would not arise unless the constraint was enforced.

To examine sequence diversity of the designed promoters, global alignment distances were determined for each pairwise combination of sequences within each promoter set. The distribution of these pairwise alignment distances was used as a proxy for sequence diversity (Supplementary Fig. 19). The results of this analysis indicate that sequence diversity was similar in the training data and designed sequences for all three design objectives. Specifically, alignment scores in promoter sets generated by the screening and evolution strategies were comparable to or less than scores in corresponding training datasets. In contrast, alignment scores in promoter sets generated by the gradient ascent approach were greater than scores in corresponding training datasets (MWT, p < 10^−50^ for all). The results suggest that the screening and evolution strategies produced promoter sets with comparable diversity to sequences generated by mining the FACS-seq training data for those exhibiting high activities, while some diversity was lost when optimizing promoter sets through the gradient ascent strategy.

### 2.4 Model designs yield large sets of promoters with high activity

We used a flow cytometry assay to individually characterize a subset of promoter sequences measured in FACS-seq to determine the reliability of measurements and compare our novel promoter designs to commonly used benchmark promoters. To determine FACS-seq measurement reliability across a range of promoter activities, we selected 3 sequences for each of 8 evenly distributed FACS-seq-derived activity values, in both the uninduced and induced conditions. To better characterize sequences that were observed in an extreme bin (Bins 1, 12), for each promoter set, we selected up to 3 sequences observed only in an extreme bin in either the uninduced or induced condition. Finally, we randomly selected sequences from each promoter set that were not characterized in FACS-seq, such that 5 sequences were measured in total.

Additionally, in order to provide further characterization for promoter sets which would be of interest to the research community, we characterized ten randomly selected sequences from one designed promoter set (referred to as the “final promoter sets”) for each of our three objectives (P_GPD_-Activity, P_ZEV_ -Induced, P_ZEV_-Activation Ratio) (Supplementary Table 1, “Final Design”). Measured activities and sequences for these promoters and control promoters (described below) appear in Supplementary Tables 3 and 4. Since the sequence design strategies were able to generate sequences with high activities even under the GC constraint, we selected promoter sets that used this constraint, and since the gradient ascent approach produced less diverse promoter sequences than other approaches, we selected promoter sets that used the evolution design approach.

We characterized the final set of 145 promoter sequences using the previously described two-color reporter construct in a flow cytometry assay, successfully measuring activities of 140 of the designs. (Results for all measured sequences appear in the Supplementary Data, in the file ‘Validation/validation_output.csv’.) In addition, we characterized a set of commonly used promoters (P_GPD_, P_TEF1_, P_ADH1_, P_PGK1_, P_TPI1_, P_CYC1_) and 3 previously described ZEV promoters (P3, P4, P8)^15^ in the same assay to benchmark the model-designed promoters (Supplementary Fig. 20). The model-designed ZEV promoters and controls were characterized in the presence and absence of 1 μM beta-estradiol. The activities of the model-designed promoters as measured in FACS-seq and individually via flow cytometry were compared (Fig. 5A). Lack of agreement in activity measurements between the two experiments was observed for sequences in which the FACS-seq measured activity was at the extremes of detection (“Offscale” sequences, Fig. 5A). In the uninduced condition, many of the P_ZEV_ sequences fell into the lower limit of detection in the FACS-seq experiment, but exhibited high activities in individual flow cytometry characterization. The results suggest that these sequences are functional, but failed to exhibit activity in FACS-seq assay (potentially due to a mutation in the expression construct outside the region covered by the sequencing). The activities measured from FACS-seq for sequences that did not fall within the limits of detection of that experiment correlated well with activities measured via flow cytometry (R^2^ = 0.92).

**Figure 5:**
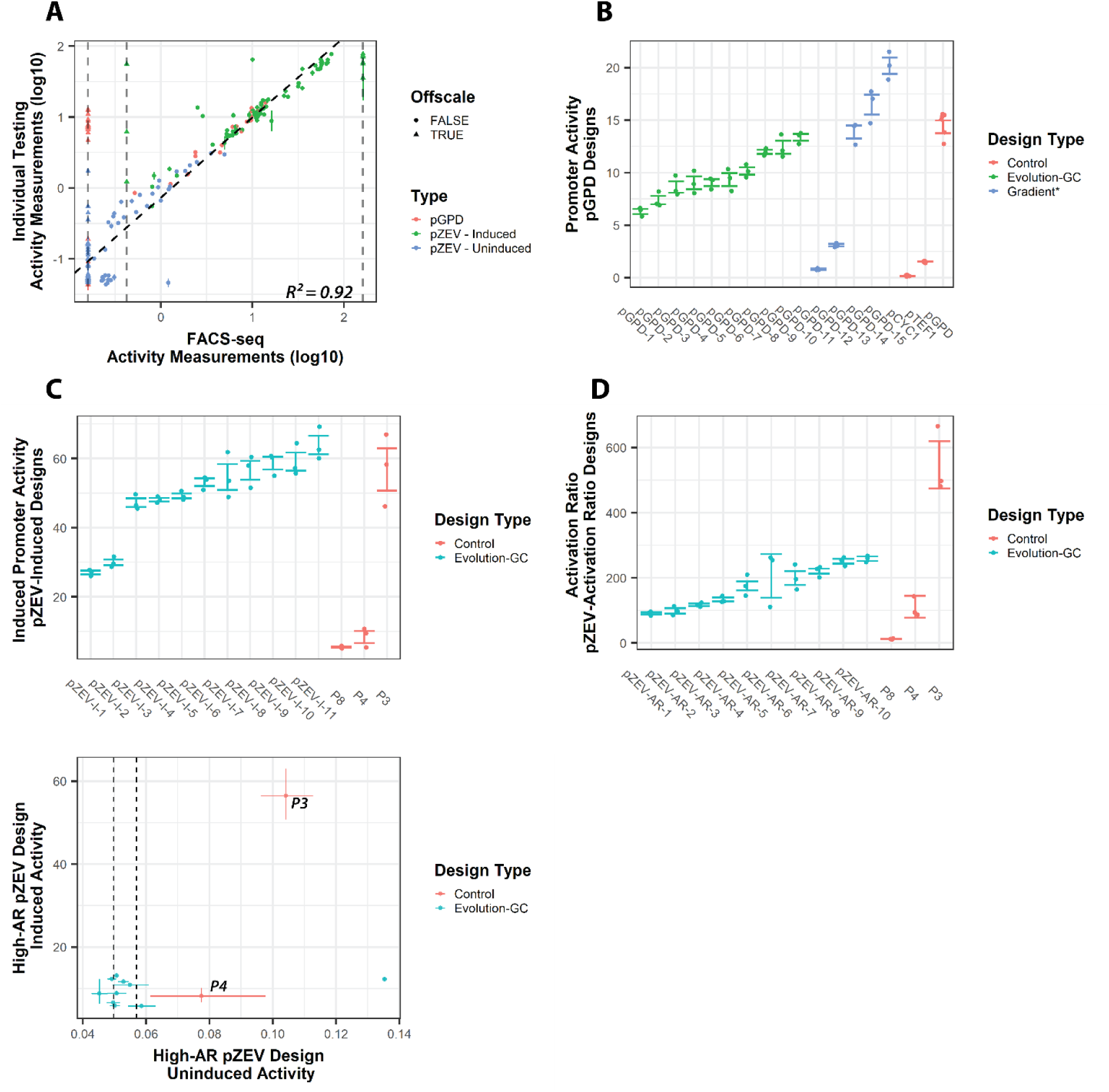
Validating activities of individual designed promoters by flow cytometry characterization. **A:** Promoter activity measurements (as base-10 logarithms) for selected sequences measured both in FACS-seq and by individual flow cytometry. “FACS-seq”: promoter activities determined from FACS-seq; “Individual Testing”: promoter activities as determined by flow cytometry (three biological replicates per condition, measuring the median ratio of fluorescence from GFP driven by the promoter of interest to mCherry driven by P_TEF1_.). Offscale sequences are those sequences which were measured in either only the lowest-activity or only the highest-activity bin during FACS-seq. Coefficient of determination (R^2^) excludes offscale sequences. **B:** Individually measured promoter activities (linear scale) determined by flow cytometry for selected P_GPD_ designs and for control sequences (“Control”). “Evolution-GC”: randomly chosen sequences from the selected P_GPD_ promoter set designed using the evolution strategy and the GC constraint; “Gradient*”: randomly chosen sequences from the promoter set designed using the gradient strategy, with an elevated threshold for selection. Control sequence names are indicated by text labels. **C:** Individually measured promoter activities (linear scale) determined by flow cytometry for a selected P_ZEV_-Induced design and for control sequences (“Control”). “Evolution-GC”: randomly chosen sequences from the selected P_ZEV_-Induced promoter set designed using the evolution strategy and the GC constraint. **D:** Individually measured promoter activities (linear scale) determined by flow cytometry for a selected P_ZEV_-Activation Ratio design and for control sequences (“Control”). “Evolution-GC”: randomly chosen sequences from the selected P_ZEV_-Activation Ratio promoter set designed using the evolution strategy and the GC constraint. **E:** Promoter activities (linear scale) in the uninduced and induced condition measured by flow cytometry for sequences appearing in **D**. Vertical dashed lines indicate the range of three independent measurements of a control plasmid (pCS4306) expressing mCherry, but not GFP. Data information: Error bars represent the standard error of three biological replicates. In panels B-D, promoter names, from left to right, are as in Supplementary Tables 3 and 4.

We next examined the results from flow cytometry validation for the final designed promoter sets. For the objective of maximizing P_GPD_ activity, in addition to the 10 sequences selected from the final P_GPD_ promoter set, we tested 5 sequences from 3 design strategies we were unable to test via FACS-seq (sets 22, 24, and 28 in Supplementary Table 1) to benchmark to the control promoter set (Fig. 5B, Supplementary Fig. 21). All tests were carried out using the mean of three biological replicates. The lowest-activity P_GPD_ sequences from our selected set had a mean activity of 6.29 (95% confidence interval (CI) [5.31, 7.45]), whereas the highest activity sequence had a mean activity of 13.36 (CI [12.06, 14.79]). By comparison, the activities of the control promoters P_TEF1_ and P_GPD_ were 1.52 (CI [1.45, 1.59]) and 14.65 (CI [13.46, 15.94]), respectively, in this same assay. Of the P_GPD_ design strategies not tested previously in FACS-seq, two (the evolution strategy, applying the GC constraint but not the extrapolation penalty; gradient ascent strategy, not applying the GC constraint and extrapolation penalty) produced similar results to our chosen final strategy (Supplementary Fig. 21). However, of the five sequences synthesized from a promoter set which used gradient ascent to reach an elevated design threshold (set 28, Supplementary Table 3), one exhibited an activity of 20.17 (CI [17.12, 23.78]) – higher than that of the P_GPD_ benchmark (p =7.27×10^−4^, one-sided *t*-test). Two sequences exhibited activities similar to P_GPD_, and two exhibited lower activities than P_GPD_ (0.80 (CI [0.60, 1.06]), 3.09 (CI [2.66, 3.59]); p < 10^−4^ for both, one-sided *t*-test). The results indicate that the gradient ascent strategy with the elevated threshold can produce promoters that exhibit higher activities than current best-in-class promoters, but occasionally incorrectly predict promoter activity, likely due to aggressive extrapolation.

We then examined how our selected final P_ZEV_-Induced designs performed relative to the benchmark ZEV promoters in the flow cytometry assay (Fig. 5C). The model-designed sequences characterized in the assay exhibited activities ranging from 27.03 (CI [24.77, 29.51]) to 63.85 (CI [53.30, 76.50]). The results demonstrate that all the tested sequences exhibited significantly higher activities than P4 and P8 (p < 0.05 for all, one-sided *t*-test), and most sequences exhibited activities comparable to P3 (56.47 (CI [35.45, 89.95]). The results showed that we were able to reliably generate ZEV promoters with induced activities comparable to the best reported sequences.

Finally, we characterized sequences from our selected final P_ZEV_-Activation Ratio design strategy in the flow cytometry assay (Fig. 5D). The measured activation ratios of the model-designed sequences ranged from 90.75 (CI [76.37, 106.37]) to 258.73 (CI [235.88, 281.71]). In contrast, the benchmark control sequences exhibited a broad range of activation ratios spanning 542.20 (CI [348.14, 815.16]) for P3, 105.94 (CI [54.57, 195.10]) for P4, and 12.62 (CI [9.97, 15.67]) for P8. In directly comparing induced and uninduced promoter activities for the model-designed sequences, we observed that induced activities for the designed sequences were comparable to that of P4 and much less than that of P3 (Fig. 5E; p < 0.01 for all, one-sided *t*-test). By contrast, uninduced activities for all but one of the designed sequences were significantly less than that of P3 (p < 0.01 for all, one-sided *t*-test), generally less than that of P4, and fell within the range of fluorescence levels observed from the no-GFP control plasmid (pCS4306) measured in the same assay. The results indicate that while the model was successful in generating a large set of sequences that were highly inducible (over 100-fold) in absolute terms, the sequences did not surpass the activation ratio of the benchmark control sequence, because uninduced activities for these sequences were too low to be accurately measured.

### 2.5 *In silico* mutagenesis elucidates design strategies driving predicted activity for novel promoters

Finally, we analyzed the model’s predictions to identify sequence features in the designed promoters that contributed to promoter activity. The analysis may shed light on our model’s approach to sequence design, as well as potentially reveal design strategies for future promoter design efforts. We applied an *in silico* mutagenesis approach, using the model’s predictions of promoter activity for single and double mutants of a designed sequence in comparison to its predictions for the original design to identify motifs of interest.

We examined all possible single mutants of a representative P_GPD_ design chosen from a promoter set designed using the screening strategy without the extrapolation penalty (Fig. 6A). Our analysis identified a sequence within the upstream spacer sequence, 5’ to the designed TFBSes, which produced sequences with substantially weaker predicted activities when mutated (boxed in Fig. 6A). We determined that the motif ‘TGASTCA’, a known binding site for the transcriptional activator Gcn4p^46^, was preferred in this site. This result suggests that our model has the capacity to identify activating binding sites for native transcription factors.

**Figure 6:**
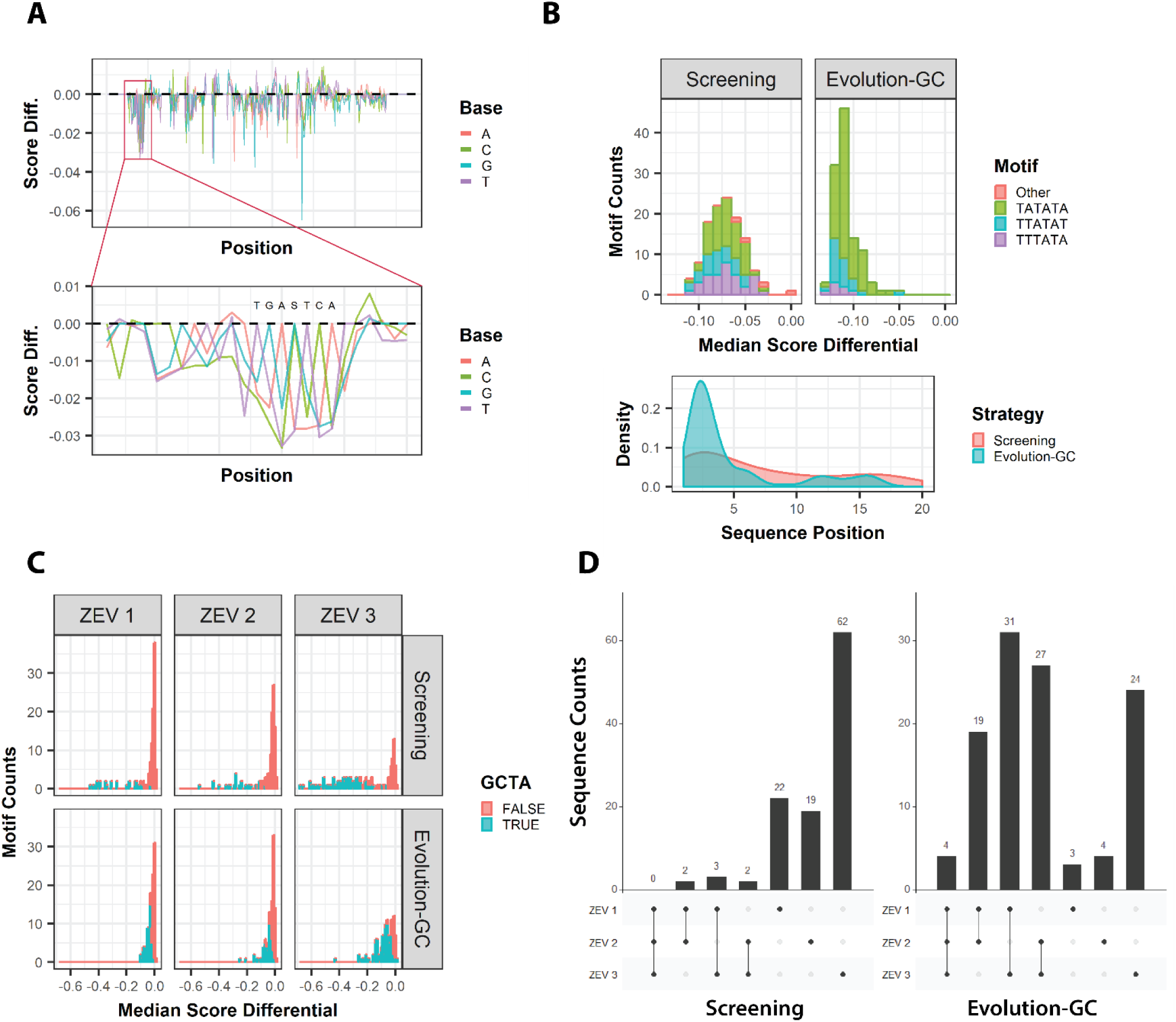
*In silico* mutagenesis enables identification of functional motifs in designed sequences. **A:** Putative motif identification via analysis of *in silico* predicted activities ofmutants in a P_GPD_ design. *Above*: predicted promoter activities of all single mutants in spacer sequences in this promoter. *Boxed region*: predicted Gcn4p site. “Score Diff.”: Score differential of each single mutant (difference between predicted activity of the mutant and predicted activity of the original sequence). *Below*: view of the boxed region, showing how mutations away from the ‘TGASTCA’ Gcn4p consensus motif are predicted to weaken the promoter. **B:** Analysis of the sequences and positions of the strongest hexamer motifs in the upstream regions of P_ZEV_ designs. *Above:* histogram of median score differential (median activity of mutants in the motif region minus original activity) for the P_ZEV_-Induced design generated using the screening strategy and the extrapolation penalty (“Screening”), and the P_ZEV_-Induced design selected for characterization of individual sequences (“Evolution-GC”). *Color*: sequence of strongest hexamer motif. *Below:* density plot of strongest motif position in the upstream regions of these sequence sets. Sequence position is numbered starting at the beginning of the designed sequence. **C:** Predicted effect of mutagenesis in the four bases immediately following ZEV ATF binding sites in designed P_ZEV_-Activation Ratio promoter sets. Histograms show median score differential of all single mutants in the four bases immediately following each ZEV ATF site (“ZEV 1”, “ZEV 2”, “ZEV 3”); color-coding indicates whether the mutagenized four-base sequence is ‘GCTA’. Sequence sets analyzed are (*above*) a P_ZEV_-Activation Ratio set generated by the screening strategy and using the extrapolation penalty (“Screening”) and (*below*) a P_ZEV_-Activation Ratio set generated by the evolution strategy and using the GC constraint and the extrapolation penalty (“Evolution-GC”). **D:** Analysis of the number of times the ‘GCTA’ motif occurs following each ZEV ATF binding site in the “Screening” and “Evolution-GC” promoter sets analyzed in **C**.

To further explore the potential of *in silico* mutageneic screening, we extended our method to include all possible double mutants of a sequence of interest. For each possible pair of mutagenized positions in a sequence, we determined the difference between predicted sequence activity for the original sequence and each single and double mutant, which we refer to as the score differential. We assumed that in the absence of an interaction between the two bases, the score differential of any double mutant would be the sum of the score differentials of the corresponding single mutants. For each double mutant, we determined the predicted score differential under this assumption, giving us a vector of expected double mutant score differentials and a vector of corresponding actual score differentials. We took the Euclidean distance between these vectors as a measure of interaction between those sequence positions, resulting in a two-dimensional grid of score differentials between each pair of sequence positions, which can be used to identify interacting bases. Applying this to the P_GPD_ design analyzed in Fig. 6A, we indeed observed interactions in the region corresponding to the putative Gcn4p site (Supplementary Fig. 22). These result suggest that our model has the capacity to identify activating binding sites for native transcription factors.

We then used the *in silico* mutagenesis approach to identify commonly used motifs in the model-guided design of entire promoter sets (Fig. 6B). We focused on a P_ZEV_-Induced sequence set generated by the screening strategy using the extrapolation penalty (“Screening”, Fig. 6B) and on the final P_ZEV_-Induced set we previously characterized by individual flow cytometry (“Evolution-GC”, Fig. 6B). Because generating activity measurements for all possible double mutants is computationally intensive, we generated only single mutants for this analysis. Strong motifs were generally located in the upstream spacer sequence of examined designs. We identified the strongest hexameric motif in the upstream spacer of each sequence in these promoter sets by defining the strength of a motif as the mean score differential of all single mutants of that motif versus the parent sequence. In almost all cases, the strongest motif was identified as ‘TATATA’, ‘TTATAT’, or ‘TTTATA’. The median change in predicted score due to a mutation in a sequence’s strongest motif was -0.07 for the Screening designs and -0.11 for the Evolution-GC designs (Fig. 6B, upper panel). We examined the position of the motifs within the designed sequences (Fig. 6B, lower panel) and found them to be dispersed throughout the upstream spacer sequence in the Screening designs, but preferentially located near the beginning of the spacer in the Evolution-GC designs. The results indicate that the model “prefers” a TATA-like motif in the upstream spacer of P_ZEV_-Induced designs, preferably near the 5’ end of the sequence. The evolution strategy enables the model to generate such sequences efficiently and place the motif in a preferred location, whereas the screening strategy selects TATA-like motifs at any point in the spacer. As this motif resembles the TATA initiation motif, we speculated that the model may have learned to place this motif to serve as an additional site for recruiting RNA polymerase. This is a surprising result, as the TATA motif is located 3’ to TFBSes in natural promoters.

We used a similar approach to investigate design strategies for promoter sets designed to maximize activation ratio, focusing on a P_ZEV_-Activation Ratio set generated by the screening strategy using the extrapolation penalty (“Screening”, Fig. 6C and 6D) and on a P_ZEV_-Activation Ratio set generated by the evolution strategy using the GC constraint and extrapolation penalty (“Evolution-GC”, Fig. 6C and 6D). We observed that in these sequences, the ZEV ATF’s binding motif, ‘GCGTGGGCG’, was frequently extended by the sequence ‘GCTA’. For each sequence in the two P_ZEV_-Activation Ratio sets examined, we determined whether the tetramer following each of the three designed ZEV ATF binding sites was ‘GCTA’, as well as the score differential in mutants of this tetramer. We found that non-GCTA motifs had a small influence on predicted activation ratio – examining Screening and Evolution-GC designs separately at all three ZEV ATF sites, the median score differential was between -0.02 and 0 at all sites. By contrast, the median score differential for Screening designs containing a GCTA motif was significantly greater at all positions (p < 10^−12^ for all, MWT): -0.27, -0.28, -0.35 at sites 1, 2, 3. The trend was less pronounced for Evolution-GC designs containing a GCTA motif (-0.04, - 0.04, -0.07 at site 1, 2, 3), although statistically significant in comparison to results for non-GCTA motifs (p < 10^−8^ for all, MWT).

We then determined the pattern of occurrence of the ‘GCTA’ motif extension in the P_ZEV_-Activation Ratio designs (Fig. 6D). We observed that double or triple occurrences of the motif were rare for Screening designs (7 of 118 sequences), but more common for Evolution-GC designs (81 of 114 sequences). This result suggests that the evolution approach was able to generate the GCTA motif extension more efficiently than the random screening approach. The result also suggests an explanation for the greater impact of GCTA mutations in the Screening set as compared to the Evolution-GC set – when multiple copies of the GCTA motif are present, at least one copy of the motif is still present even if a mutation disrupts another copy, which could reduce the predicted impact of disrupting one copy of the motif. Supporting this explanation, the median score differential for GCTA mutations in Evolution-GC sequences with only one GCTA motif was significantly greater than for sequences with multiple copies of the motif (p =1.88×10^−10^, MWT) (Supplementary Fig. 23).

## 3. Discussion

We developed a CNN model which accurately predicts promoter activity in two artificial yeast promoter libraries, and developed design strategies exploiting this model to generate large sets of sequence-diverse promoters. Our modeling approach builds on previous work modeling relatively short sequence libraries in the 50-bp length range^27^. While most previous efforts have been restricted to modeling only the 5’ UTR of an otherwise constant promoter, we measured activities for full-length promoters by developing a novel FACS-seq pipeline that integrates data collected from two NGS platforms. We used a low-read, full-length sequencing run to determine the sequence of each variant, and a high-read run sequencing only part of each variant to determine the variant’s abundance in each bin. Our model’s ability to predict promoter activity for a complex sequence containing many different elements (e.g., TFBSes and other conserved motifs, core promoter, 5’ UTR) exemplifies the ability of deep neural networks to model complex data.

We tested a variety of sequence design approaches, and found that the properties of the resulting sequences varied substantially from approach to approach. Sequences generated by the screening strategy did not outperform sequences with the highest activities measured in the original library datasets. However, when optimizing for P_GPD_ activity and P_ZEV_-induced activity, the evolution and gradient ascent strategies generated sequences with activities comparable to or greater than benchmark promoters. In addition, we generated high-performing sequences even when applying constraints to GC content. These results demonstrate the usefulness of model-guided design for producing sequences with useful, rare properties.

P_ZEV_-Activation Ratio designs had lower apparent activation ratios than the benchmark P3 control promoter due to challenges in measuring very low promoter activities accurately.

Measured activities in the uninduced condition for P_ZEV_-Activation Ratio designs were at the lower limit of detection, while P3 had a measurable level of uninduced expression (Fig. 5E). While activation ratio is an intuitive and simple measure of promoter inducibility, it has limitations when applied to sequences that have very low uninduced activities. Developing a design strategy that independently optimizes for high induced and low uninduced activity may address the limitations of using activation ratios as a metric of inducibility.

Beyond generating large promoter sets, we used the model to explore strategies that yielded promoters with high predicted activity or activation ratio. The model’s ability to identify significant motifs in an unbiased manner (Fig. 6A) suggests that our approach may be valuable for future studies of native yeast promoter regulation. Unexpectedly, the model placed TATA-like motifs at the 5’ end of P_ZEV_-Induced designed promoters. Further characterization is needed to directly determine the functional role of the TATA-like motifs, but this “design choice” may be of interest to future rational promoter design efforts. We additionally found that P_ZEV_-Activation Ratio designed promoters often contained an apparent four-base extension (‘GCTA’) of the ZEV ATF binding site. Although further characterization is needed, it is possible that this extension acts by increasing the sequence’s binding affinity for the ZEV ATF, thus increasing ZEV ATF-dependent transcription. This hypothesis is supported by previous studies demonstrating a role for bases outside transcription factors’ “core motifs” in determining their binding affinity to DNA sequences^47^, including for the Zif-268 transcription factor used in ZEV ATF^48^. Thus, our modeling approach could be applied to better characterize transcription factor-DNA sequence binding affinities.

By generating large promoter sets with activities comparable to or outperforming state-of-the-art benchmark promoter sequences, our work creates a useful tool for applications in metabolic engineering and synthetic biology. The approaches used in our work are not necessarily limited to yeast or to promoter elements, and could be applied to expression parts for use in other organisms, such as mammalian cell lines, or to designing other classes of DNA sequences, such as terminators or RNA switches. Our results demonstrate that high-throughput characterization of artificial DNA sequence libraries enables accurate modeling of the DNA sequence-function relationship, which in turn enables the design of novel DNA sequences fulfilling specified design constraints.

## 4. Materials and Methods

Plasmids used in this study are described in Supplementary Table 5, and yeast strains used in this study are described in Supplementary Table 6. Expand High Fidelity PCR system (Roche Diagnostics) was used for PCR amplifications according to manufacturer’s instructions, unless described otherwise. Sanger sequencing was performed by Elim Biopharmaceuticals, Inc. Oligonucleotides used in generating plasmids and yeast strains appear in Supplementary Table 7.

### 4.1 Random library design and assembly

Libraries were assembled by PCR-amplifying oligonucleotides (Stanford School of Medicine Protein & Nucleic Acid Facility (PAN); Integrated DNA Technologies) corresponding to each spacer sequence, joining them via Golden Gate assembly^49^ using constant sites located within constant regions, and PCR-amplifying the resulting products with primers providing 40 bp of homology to vector plasmids on each side. Oligonucleotides and PCR primers used to assemble libraries are listed in Supplementary Table 8. The combinations of oligonucleotides used to PCR-amplify each fragment are given in Supplementary Table 9. PCR and Golden Gate reaction conditions are given in Supplementary Note 1; briefly, 15 fmol of each DNA fragment was used in 10-µL, 50-cycle Golden Gate reactions.

### 4.2 Yeast strains, culture, transformation, and passaging for FACS-seq

*Saccharomyces cerevisiae* strain CSY3 (W303 MATα) was used in the P_GPD_ library experiment. To create a strain expressing the ZEV artificial transcription factor for the P_ZEV_ library experiment, the P_ACT1_ promoter and the ZEV artificial transcription factor gene were PCR-amplified in a single fragment from DBY19053^15^, and a fragment containing the T_CYC1_ terminator and the entire plasmid backbone was PCR-amplified from pCS2657^50^. These fragments were joined by Gibson assembly^51^, yielding pCS4339; the P_ACT1_ – ZEV ATF – T_CYC1_ expression cassette was PCR-amplified and integrated into the LEU2 locus of CSY3 using the Cas9-assisted integration method^52^ (using pCS4187^53^ as the guide RNA plasmid), yielding strain CSY1252. CSY1252 was also used in the promoter design validation experiments.

The plasmid pCS1748^39^ expresses GFP and mCherry from separate copies of the P_TEF1_ promoter. The plasmid pCS4305 was generated by digesting pCS1748 with ClaI and MfeI to remove the P_TEF1_ driving GFP expression; a 675-bp sequence encoding P_GPD_ and part of the 3’ UTR of YGR193C (Supplementary Note 2), and a sequence replacing a portion of GFP deleted by restriction digestion, were inserted using Gibson assembly^51^. The P_GPD_ sequence was PCR-amplified from pCS2656^50^, with bases added at the 3’ end to match the 5’ UTR of the S288C reference wild-type P_GPD_ sequence^54^. To generate a vector for library integration, this plasmid was further modified by removing the P_GPD_ promoter by digestion with ClaI and MfeI and using Gibson assembly to insert a sequence beginning with the first 60 bp of the YGR193C 3’ UTR sequence present in the original P_GPD_ promoter (to ensure that all promoters were tested in a consistent genetic context). A ZraI cut site was created by adding a ‘C’ nucleotide at the end of this 60-bp sequence, and the excised yEGFP sequence, lacking the first two bases of the yEGFP start codon, followed, yielding pCS4306 (Supplementary Fig. 24). To clone a promoter library into yeast, pCS4306 was linearized with ZraI digestion, and yeast were co-transformed with linearized plasmid and the library insert. This co-transformation was carried out as previously described^20^. Briefly, 50 ml yeast culture (OD_600_ 1.3–1.5) was incubated with Tris-DTT buffer (2.5 M DTT, 1 M Tris, pH 8.0) for 15-20 min at 30°C, pelleted, washed, and resuspended in Buffer E (10 mM Tris, pH 7.5, 2 mM MgCl_2_) to 200 µl. To 50 µl of the yeast cell suspension, 2 µg of linearized plasmid and 1 µg of library insert DNA was added and the DNA-cell suspension was electroporated (2 mm gap cuvette, 540 V, 25 µF, infinite resistance). Transformed cells were diluted to 1 ml volume in yeast peptone dextrose (YPD) media, incubated for 1 hour, then further diluted in yeast nitrogen base medium (BD Diagnostics) lacking uracil and containing 2% dextrose (YNB-U).

All libraries were grown in YNB-U, and passaged at least three times before sorting, with at least 10 OD_600_*mL units transferred in each passage. For experiments involving P_ZEV_ promoters, separate cultures with and without 1 µM beta-estradiol added were started 18 hours before the sort. Cultures were back-diluted to an OD of 0.05-0.1 5 hours before the sort to maintain them in log phase.

### 4.3 Library sorting

Cultures were harvested at an OD_600_ of 0.7-0.8, spun down, and resuspended in phosphate-buffered saline (PBS) with 10 µg/mL DAPI (ThermoFisher). The sorts were performed on a FACSAria II cell sorter (BD Biosciences) with excitation and emission filters for GFP, mCherry, and DAPI as previously described^20^. Briefly, GFP was excited at 488 nm and measured with a splitter of 505 nm and bandpass filter of 525/50 nm, mCherry was excited at 532 nm and measured with a splitter of 600 nm and bandpass filter of 610/20 nm, and DAPI was excited at 355 nm and measured with a bandpass filter of 450/50 nm. Viable cells (as identified by a viability gate based on DAPI fluorescence and side-scatter area) were sorted into one of twelve bins of equal width on the basis of the GFP/mCherry ratio. These bins were chosen to cover the range of promoter activities present in the sorted library. The sort gates were generated using a MATLAB script.

Four bins were collected at a time, in three passes: one collecting bins 1, 4, 7, and 10, one collecting bins 2, 5, 8, and 11, and one collecting bins 3, 6, 9, and 12. For each pass, cells were collected until a target number of viable cells had been sorted. In the P_GPD_ experiment, two replicates were collected. In experiments involving P_ZEV_ inducible promoters, the 12 bins were each collected once for the uninduced and once for the induced condition. In these experiments, the gating and cytometry parameters were set separately for the uninduced and induced conditions. Counts of cells sorted per bin for the P_GPD_ experiment are in Supplementary Table 10, for P_ZEV_ in Supplementary Table 11, and for the validation FACS-seq experiment testing designed promoters in Supplementary Table 12.

The sort parameters used for the P_GPD_ experiment were treated as reference conditions for experiments involving inducible promoters. To relate measurements from experiments involving inducible promoters to the results of the P_GPD_ experiment, flow cytometry data collected for the libraries under the conditions used for sorting and under the reference conditions (those used to sort the P_GPD_ library) was used as a benchmark to convert the GFP/mCherry ratios used as bin edges to their equivalents under the parameters used for the P_GPD_ sort (Supplementary Fig. 25). Promoter activities were calculated for each cell, and approximately corresponding cells in each sample were identified by sorting these values. A linear model was fit, and bin edges used in the experiment as measured were converted to their equivalents under the reference conditions. For the validation FACS-seq, the fit was carried out using the mean of three samples collected under the experimental conditions and compared to one sample collected under the reference conditions.

### 4.4 NGS sample preparation

After sorting, cells were grown to saturation in YNB-U. 1.5-mL aliquots of cell culture were used as input in minipreps with the Zymoprep Yeast Plasmid Miniprep II kit (Zymo Research) according to manufacturer’s instructions. Multiple minipreps were performed where necessary so that one miniprep was performed for every 1,000,000 cells collected. In experiments involving inducible promoters, unsorted cells from the uninduced and induced libraries were also regrown and miniprepped. For each bin, the entire miniprepped volume was used as template in a PCR reaction using the KAPA HiFi PCR Kit (Kapa Biosystems). A ten-fold dilution of this PCR product was used as template in a barcoding PCR adding Illumina adapter sequences and dual barcodes. As a quality-control measure, variable-length sequences were included in each primer immediately 5’ to the sequence annealing region, serving as a backup barcoding method. These PCRs were purified using the DNA Clean & Concentrator kit (Zymo Research) according to manufacturer’ instructions, quantitated using a Qubit fluorometer (ThermoFisher), and mixed at a ratio calculated to provide an approximately equal number of sequencing reads for each cell originally collected. This mixdown was then gel-extracted on a gel containing SybrSafe Red (ThermoFisher) and 2% agarose, PCR-amplified for 5-6 cycles starting from a concentration of 1 nM with primers corresponding to Illumina adapters to ensure full-length products, and purified, yielding the final NGS samples. PCR reaction parameters are given in Supplementary Note 1, oligonucleotides used in NGS sample prep are given in Supplementary Table 13, and oligonucleotide choices for the P_GPD_, P_ZEV_, and design validation FACS-seq appear in Supplementary Tables 14, 15, and 16, respectively.

Sample quality was checked using a Bioanalyzer 2100 (Agilent). Next-generation sequencing was carried out on an Illumina MiSeq by either PAN or the Chan Zuckerberg Biohub, using 2×300 paired-end reads, with PhiX sequencing control added to 30% by molarity to increase diversity in constant or AT-rich regions. In some experiments, the sample was additionally sequenced on an Illumina NextSeq by the Biohub, using 1×75 unpaired reads.

### 4.5 NGS processing: Error-tolerant sequence determination

NGS data processing and model training were carried out on Google Cloud virtual machines, using Nvidia Tesla K80 GPUs.

Sequences in the sorted libraries were determined using the MiSeq output. Paired-end reads were first merged using Paired-End reAd mergeR (PEAR) version 0.9.6^55^. In the P_GPD_ and P_ZEV_ experiments, there was a risk that errors in PCR or mutations during passage after FACS sorting could give rise to similar sequences, differing at only a few positions. These “sibling sequences” would likely have similar activities, and could lead to inflated estimates of model quality if one appeared in training data and another in validation or test data. To avoid this risk, we chose to group reads with similar sequences together and determine a consensus sequence. We sorted the sequences and compared each sequence with the one following using Needleman-Wunsch alignment, with a gap penalty of 5 and a mismatch penalty of 1. Needleman-Wunsch alignments were carried out using a parallelized implementation of the algorithm (https://github.com/hgbrian/nwalign)^56^. For each experiment, we established a cutoff value for similarity, and used this to determine where groups of related sequences began and ended (Supplementary Fig. 26).

Because sequences are arranged in alphabetical order in this process, mutations near the start of a sequence need to be accounted for separately. To do this, we repeated the process after reversing and re-sorting the original sequences; each read was thus assigned to two clusters, one from each sorting. Clusters with reads in common were then merged to yield the final groups of reads corresponding to each sequence.

Read clusters were reduced to consensus sequences by taking a majority vote at each position in the sequence to obtain a consensus call for that position. Singleton sequences and sequences without a majority call at each position were discarded (Supplementary Fig. 27).

### 4.6 NGS processing: Measuring promoter activity

In the P_GPD_ and P_ZEV_ experiments, the MiSeq runs yielded 0.5 or fewer reads per original cell for most bins. For many sequences, there were enough reads to identify the sequence itself, but not enough to accurately quantitate promoter activity. To solve this problem, the samples were resequenced on an Illumina NextSeq using a 1×75 single read kit (for lack of paired-end kits long enough to sequence this sample in its entirety on NextSeq). This allowed many more sequences to be accurately quantitated (Supplementary Fig. 28). Measures of promoter activity derived from NextSeq data and full promoter sequences derived from MiSeq data were related using the first 35 bp of each library member, which is fully randomized and was found to act as a unique identifier for almost every sequence. In the validation experiment, the MiSeq data was used to calculate promoter activity directly, since this experiment featured a relatively small number of designed sequences.

Promoter activities were obtained following previously described methods^20, 21^. First, each read was assigned to a bin by identifying the barcoding oligos used to generate it; each oligo contained either a variable-length “skew sequence” read as part of the sequencing read or a barcode, determined in a separate barcoding read. Sequences with fewer than a threshold number of reads measured in each replicate were discarded. The choice of skew sequences and barcodes used to assign reads to bins in each experiment, and the read count thresholds used in each experiment, are provided in Supplementary Table 17. Read counts in each bin *i* were then normalized by multiplying by *C_i_/R_i_*, where *C_i_* is the number of cells collected in bin *i*, and *R_i_* is the total number of reads observed in the bin. A maximum-likelihood estimation process was used to assign a mean to each sequence. As much of this process as possible was executed in parallel on the GPU, using the Numba project’s CUDA libraries (numba.pydata.org).

When estimating means for each sequence, each replicate of each experiment was processed separately. Based on prior experience^20^ and the results of preliminary experiments, it was assumed that the distribution of fluorescence for each sequence in a library was log-normal, and that the standard deviation of the fluorescence distribution *σ* was the same for all sequences. To provide some robustness against outliers, it was further assumed that with a probability *ε*, cells were collected not from the log-normal distribution, but from a uniform distribution across all bins.

The mean estimation process then requires two hyperparameters: *σ* and *ε*. For a given promoter activity *μ*, a given *σ* and *ε*, and a number of bins *N*, the probability of a cell being observed in a bin *i* with edges *a_i_* and *b_i_* is 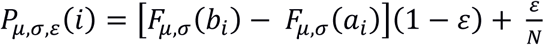, where *F_μ,σ_* is the cumulative distribution function of a normal distribution with mean *μ* and standard deviation *σ*. Supposing that the number of cells observed in bin *i* is *r_i_*, the log-likelihood of a set of parameters given observed data is 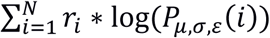^21^. Let (*σ*^*^, *ε*^*^) be a choice of values for (*σ*, *ε*). For a set of *M* possible values of *μ*, we can construct an NxM matrix *W*_(*σ*^*^,*ε*^*^)_, where *W*_(*σ*^*^,*ε*^*^)*ij*_ = log (*P*_*μ*_j_,*σ*^*^,*ε*^*^_ (*i*)). Supposing there are *S* sequences total, if the SxN matrix *A* contains the number of cells observed in each bin for each sequence, the product *AW* is an SxM matrix giving the log-likelihood of each possible value of *μ* for each sequence. Estimates of *μ* are chosen for each sequence to maximize log-likelihood, and the sum of the log-likelihoods of each sequence acts as a score for the original choice of *σ* and *ε*.

Using parallel computation on a GPU to accelerate this process enabled us to optimize the hyperparameters used in fitting via a grid search, to determine the sensitivity of fits to hyperparameter choice. Optimal hyperparameter values were found for each replicate or condition in each experiment (Supplementary Fig. 29, Supplementary Table 18). Additionally, we tested the sensitivity of the fit to the choice of hyperparameter scores, by calculating the root-mean-squared distance from the vector of fit means under the final hyperparameter values to the vector of fit means under each other choice of hyperparameters tested (Supplementary Fig. 29).

This resulted in a table of sequences and promoter activity values – for the P_GPD_ experiment, promoter activity was measured once in each replicate, while in the P_ZEV_ and the validation experiment, it was measured once in the uninduced and once in the induced condition. Sequences with mutations in the designed constant regions were removed, as well as sequences with reads in only the lowest-activity or highest-activity bin. The resulting table was then used to train models of promoter activity.

### 4.7 Model implementation and training

Models were implemented in Keras (keras.io) version 2.1.6, with Tensorflow version 1.7.0 as the backend. The architecture described above, under the heading “A convolutional neural network can accurately predict promoter activities”, was used for all models. A training/validation/test split of 80%:10%:10% was used in training the models. All sequences were padded on each side with at least 25 bp of the surrounding pCS4306 vector sequence; to account for the different lengths of P_GPD_ and P_ZEV_ promoters when training models on the merged datasets, the pads were extended for P_ZEV_ promoters, to a total length of 58 bp on each side. During training and validation, the dataset was augmented by applying a shift of 0 to 7 bp.

Models were trained with Adam as the optimizer^57^, with a learning rate of 10^−5^. Huber loss with *δ* = 0.15 was used as the loss function. When the model had two outputs, both were weighted equally in calculating the loss. Model training was terminated using early stopping after five consecutive epochs with no improvement in validation loss.

### 4.8 Designing novel promoters

The strategies tested for designing novel promoters are summarized in Supplementary Table 1. Three objectives were optimized: overall activity for P_GPD_ designs, activity under beta-estradiol induction for P_ZEV_ designs, and activation ratio for P_ZEV_ designs. In all cases, at least 100 promoters with the objective predicted to be above a design threshold were designed. The design thresholds were chosen empirically to generate the desired promoter sets without expending an unreasonable amount of computational resources.

The predictions for each sequence were generated using the set of 9 models trained on the joined datasets. Predictions from the models were merged either by taking the mean of the individual predictions, or the mean of predictions minus their standard deviation (to buffer the possible effect of outlier predictions). To increase the diversity and ease of assembly of the designed sequences, a filter was applied in some experiments to reject sequences containing regions 20 bp or longer with GC content below 25% or above 80%.

As described in ‘Results’, three design strategies were tested: screening, in silico evolution, and gradient ascent. In screening, sets of sequences were randomly generated, using the same constant regions and base composition probabilities used in designing the original libraries. These sequences were then tested and accepted if they met the objective. In in silico evolution, a set of sequences was iteratively generated from a randomly chosen parent sequence and tested; the highest-scoring sequence was passed on to the next round of evolution. The number of mutations induced in each round decreased over time. In gradient ascent, initially random sequences are iteratively modified by having the model predict what incremental change would most increase the predicted score. Values of cycle-dependent parameters used in the evolution and gradient-ascent strategies are given in Supplementary Table 19 and Supplementary Table 20, respectively. The choices of parameters used to specify each experiment (target promoter, objective to maximize, use of optional GC filter, function used to merge sub-model outputs, design strategy, final score threshold) are provided in Supplementary Table 1.

### 4.9 Assembling designed sequences from an oligo pool

New sequences derived from the sequence evolution strategies or selected from the original FACS-seq data as controls were assembled from an oligonucleotide pool (Twist Bioscience). Each sequence set was assigned unique PCR amplification sites, designed using a Python script to minimize cross-talk between pools; oligo annealing temperatures in this script were calculated using the Primer3 library^58^. Each sequence was then designed as a pair of oligos (a forward and reverse oligo), which could be joined by Golden Gate assembly using a unique assembly site.

Forward and reverse oligos for each sequence in each sequence set were amplified from the oligonucleotide pool in KAPA PCR reactions, using selective primers complementary to the designed unique PCR amplification sites to selectively amplify the desired subpool. The designs for each sequence set were then assembled by Golden Gate assembly (following the reaction conditions described in Supplementary Note 1 as “Golden Gate Assembly of Libraries”) and further PCR-amplified. An equimolar mixture of the resulting subpools was then cloned into pCS4306 by gap repair as described above.

### 4.10 Testing individual sequences

To validate FACS-seq results and further characterize novel promoters, a subset of 145 sequences was chosen to be synthesized and tested individually. Sequences were chosen using an R script to obtain a minimal set of sequences needed to test hypotheses of interest. To clone single promoters, the sequences were ordered from Twist Bioscience, or in the case of pre-existing control promoters, PCR-amplified using Expand High Fidelity PCR from plasmids: P_GPD_ and P_TEF1_ from pCS4305, P_ADH1_ from pCS2660, P_PGK1_ from pCS2663, P_TPI1_ from pCS2661, P_CYC1_ from pCS2659, P3 from pCS4307, P4 from pCS4308, and P8 from pCS4309. Plasmids pCS4307, pCS4308, and pCS4309 were constructed by digesting pCS4306 with ZraI and using Gibson assembly to insert the corresponding ZEV promoter sequence, which was amplified from gDNA of a yeast strain containing the sequence (DBY19053 for P3, DBY19059 for P8) or (in the case of P4) artificially synthesized as a gBlock Gene Fragment (IDT). We were unable to PCR-amplify the P4 sequence from gDNA, or have it synthesized as originally specified; we replaced the second of six closely spaced ‘GCGTGGGCG’ sites in the original sequence with ‘TTACTCAAG’. Sequences were cloned into pCS4306 by gap repair using the Frozen-EZ Yeast Transformation II Kit (Zymo Research) according to manufacturer’s instructions. Colonies were inoculated into 500 uL YNB-U liquid media in 96-well plates and grown with shaking at 30 C overnight; 5 uL of the resulting seed cultures were used to inoculate new 500 uL cultures. These were assayed on a MACSQuant VYB flow cytometer (Miltenyi Biotec GmbH) after 24 hours further growth. Confidence intervals were calculated using a *t*-test with the appropriate degrees of freedom for each sample. Data were confirmed to be normally distributed (conditional on the sequences tested) using a Q-Q plot (Supplementary Fig. 30).

### 4.11 Motif identification by *in silico* mutagenesis

Single and double mutants of sequences to be characterized by mutagenesis were generated, and activities and activation ratios estimated, using a Python script. We used a Python script to calculate score differentials between activity predictions for double mutants and single mutants, as described above under the heading “*In silico* mutagenesis elucidates ‘design strategies’ driving predicted activity for novel promoters,” as well as to identify strong motifs in the upstream spacers of P_ZEV_-Induced and P_ZEV_-Activation Ratio designed promoters.

## Data and Software Availability

The datasets and computer code produced in this study are available in the following databases:

- NGS data: Gene Expression Omnibus GSE135464 (https://www.ncbi.nlm.nih.gov/geo/query/acc.cgi?acc=GSE135464; reviewer access token: kjevqsewdvmphup).
- Other data: Zenodo (https://zenodo.org/record/3376951).
- Data analysis code: Github (https://github.com/smolkelab/promoter_design).

## Acknowledgements

We thank Deze Kong, Peter Dykstra, Aaron Cravens, and Prashanth Srinivasan for valuable comments on the manuscript. We thank Aaron Cravens for assistance with FACS sorts and for providing Cas9 plasmid pCS4187, and Travis Horst for assisting in the construction of strain CSY1252 and plasmids pCS4307-9 and pCS4339. Yeast strains DBY19053, DBY19054, and DBY19059, containing components of the ZEV system, were generously provided by Dr. David Botstein and Dr. Patrick Gibney. This work was supported by the National Institutes of Health (grants to C.D.S.) and Chan-Zuckerberg Biohub (award to C.D.S), as well as the Stanford Bio-X Institute and the Siebel Scholars Foundation (graduate student fellowships to B.J.K.).

## Author contributions

B.J.K. and C.D.S. designed the project and wrote the article. B.J.K. performed experiments and conducted data analysis.

## References

1. Ghodasara, A. & Voigt, C. A. Balancing gene expression without library construction via a reusable sRNA pool. Nucleic Acids Res. 45, 8116–8127 (2017).

2. Lee, M. E., Aswani, A., Han, A. S., Tomlin, C. J. & Dueber, J. E. Expression-level optimization of a multi-enzyme pathway in the absence of a high-throughput assay. Nucleic Acids Res. 41, 10668–78 (2013).

3. Pitera, D. J., Paddon, C. J., Newman, J. D. & Keasling, J. D. Balancing a heterologous mevalonate pathway for improved isoprenoid production in Escherichia coli. Metab. Eng. 9, 193–207 (2007).

4. Nielsen, A. A. K. et al. Genetic circuit design automation. Science 352, aac7341 (2016).

5. Rantasalo, A., Kuivanen, J., Penttilä, M., Jäntti, J. & Mojzita, D. Synthetic Toolkit for Complex Genetic Circuit Engineering in Saccharomyces cerevisiae. ACS Synth. Biol. 7, 1573–1587 (2018).

6. Shao, Z., Zhao, H. & Zhao, H. DNA assembler, an in vivo genetic method for rapid construction of biochemical pathways. Nucleic Acids Res. 37, e16 (2009).

7. Galanie, S., Thodey, K., Trenchard, I. J., Filsinger Interrante, M. & Smolke, C. D. Complete biosynthesis of opioids in yeast. Science 349, 1095–100 (2015).

8. Brown, S., Clastre, M., Courdavault, V. & O’Connor, S. E. De novo production of the plant-derived alkaloid strictosidine in yeast. Proc. Natl. Acad. Sci. U. S. A. 112, 3205–10 (2015).

9. Richardson, S. M. et al. Design of a synthetic yeast genome. Science 355, 1040–1044 (2017).

10. Gander, M. W., Vrana, J. D., Voje, W. E., Carothers, J. M. & Klavins, E. Digital logic circuits in yeast with CRISPR-dCas9 NOR gates. Nat. Commun. 8, 15459 (2017).

11. Harvey, C. J. B. et al. HEx: A heterologous expression platform for the discovery of fungal natural products. Sci. Adv. 4, eaar5459 (2018).

12. Redden, H. & Alper, H. S. The development and characterization of synthetic minimal yeast promoters. Nat. Commun. 6, 7810 (2015).

13. Alper, H., Fischer, C., Nevoigt, E. & Stephanopoulos, G. Tuning genetic control through promoter engineering. Proc. Natl. Acad. Sci. U. S. A. 102, 12678–83 (2005).

14. Blount, B. A., Weenink, T., Vasylechko, S. & Ellis, T. Rational diversification of a promoter providing fine-tuned expression and orthogonal regulation for synthetic biology. PLoS One 7, e33279 (2012).

15. McIsaac, R. S., Gibney, P. A., Chandran, S. S., Benjamin, K. R. & Botstein, D. Synthetic biology tools for programming gene expression without nutritional perturbations in Saccharomyces cerevisiae. Nucleic Acids Res. 42, e48 (2014).

16. Kolodner, R. D., Putnam, C. D. & Myung, K. Maintenance of genome stability in Saccharomyces cerevisiae. Science 297, 552–7 (2002).

17. Xi, L. et al. Predicting nucleosome positioning using a duration Hidden Markov Model. BMC Bioinformatics 11, 346 (2010).

18. Field, Y. et al. Distinct modes of regulation by chromatin encoded through nucleosome positioning signals. PLoS Comput. Biol. 4, e1000216 (2008).

19. Curran, K. A. et al. Design of synthetic yeast promoters via tuning of nucleosome architecture. Nat. Commun. 5, 4002 (2014).

20. Townshend, B., Kennedy, A. B., Xiang, J. S. & Smolke, C. D. High-throughput cellular RNA device engineering. Nat. Methods (2015). doi:10.1038/nmeth.3486

21. Peterman, N. & Levine, E. Sort-seq under the hood: implications of design choices on large-scale characterization of sequence-function relations. BMC Genomics 17, 206 (2016).

22. Dvir, S. et al. Deciphering the rules by which 5’-UTR sequences affect protein expression in yeast. Proc. Natl. Acad. Sci. U. S. A. 110, E2792–801 (2013).

23. Sharon, E. et al. Inferring gene regulatory logic from high-throughput measurements of thousands of systematically designed promoters. Nat. Biotechnol. 30, 521–30 (2012).

24. Lubliner, S. et al. Core promoter sequence in yeast is a major determinant of expression level. Genome Res. 25, 1008–17 (2015).

25. Alipanahi, B., Delong, A., Weirauch, M. T. & Frey, B. J. Predicting the sequence specificities of DNA- and RNA-binding proteins by deep learning. Nat. Biotechnol. 33, 831–8 (2015).

26. Kelley, D. R., Snoek, J. & Rinn, J. L. Basset: learning the regulatory code of the accessible genome with deep convolutional neural networks. Genome Res. 26, 990–9 (2016).

27. Cuperus, J. T. et al. Deep learning of the regulatory grammar of yeast 5’ untranslated regions from 500,000 random sequences. Genome Res. 27, 2015–2024 (2017).

28. Sample, P. et al. supplementary materials human 5’UTR design and variant effect prediction from a massively parallel translation assay. bioRxiv 1–10 (2018).

29. Da Silva, N. A. & Srikrishnan, S. Introduction and expression of genes for metabolic engineering applications in Saccharomyces cerevisiae. FEMS Yeast Res. 12, 197–214 (2012).

30. Blazeck, J., Garg, R., Reed, B. & Alper, H. S. Controlling promoter strength and regulation in Saccharomyces cerevisiae using synthetic hybrid promoters. Biotechnol. Bioeng. 109, 2884–95 (2012).

31. Hahn, S. & Young, E. T. Transcriptional regulation in Saccharomyces cerevisiae: transcription factor regulation and function, mechanisms of initiation, and roles of activators and coactivators. Genetics 189, 705–36 (2011).

32. Rojas-Duran, M. F. & Gilbert, W. V. Alternative transcription start site selection leads to large differences in translation activity in yeast. RNA 18, 2299–305 (2012).

33. Kuehner, J. N. & Brow, D. A. Quantitative analysis of in vivo initiator selection by yeast RNA polymerase II supports a scanning model. J. Biol. Chem. 281, 14119–28 (2006).

34. Kostrewa, D. et al. RNA polymerase II-TFIIB structure and mechanism of transcription initiation. Nature 462, 323–30 (2009).

35. Lubliner, S., Keren, L. & Segal, E. Sequence features of yeast and human core promoters that are predictive of maximal promoter activity. Nucleic Acids Res. 41, 5569–81 (2013).

36. Hinnebusch, A. G., Ivanov, I. P. & Sonenberg, N. Translational control by 5’-untranslated regions of eukaryotic mRNAs. Science 352, 1413–6 (2016).

37. Bitter, G. A., Chang, K. K. & Egan, K. M. A multi-component upstream activation sequence of the Saccharomyces cerevisiae glyceraldehyde-3-phosphate dehydrogenase gene promoter. Mol. Gen. Genet. 231, 22–32 (1991).

38. Khan, A., et al. JASPAR 2018: update of the open-access database of transcription factor binding profiles and its web framework. Nucleic Acids Res. 46, D260–D266 (2018).

39. Liang, J. C., Chang, A. L., Kennedy, A. B. & Smolke, C. D. A high-throughput, quantitative cell-based screen for efficient tailoring of RNA device activity. Nucleic Acids Res. 40, e154 (2012).

40. McIsaac, R. S. et al. Synthetic gene expression perturbation systems with rapid, tunable, single-gene specificity in yeast. Nucleic Acids Res. 41, e57 (2013).

41. Zou, J. et al. A primer on deep learning in genomics. Nat. Genet. (2018). doi:10.1038/s41588-018-0295-5

42. Angermueller, C., Pärnamaa, T., Parts, L. & Stegle, O. Deep learning for computational biology. Mol. Syst. Biol. 12, 878 (2016).

43. Simonyan, K., Vedaldi, A. & Zisserman, A. Deep Inside Convolutional Networks: Visualising Image Classification Models and Saliency Maps. (2013).

44. Erhan, D.; Bengio, Y.; Courville, A.; Vincen, P. Visualizing higher-layer features of a deep network. Tech. Rep. 1341, Univ. Montr. (2009).

45. Aird, D. et al. Analyzing and minimizing PCR amplification bias in Illumina sequencing libraries. Genome Biol. 12, R18 (2011).

46. Teixeira, M. C. et al. YEASTRACT: an upgraded database for the analysis of transcription regulatory networks in Saccharomyces cerevisiae. Nucleic Acids Res. 46, D348–D353 (2018).

47. Levo, M. et al. Unraveling determinants of transcription factor binding outside the core binding site. Genome Res. 25, 1018–29 (2015).

48. Rudnizky, S. et al. Single-molecule DNA unzipping reveals asymmetric modulation of a transcription factor by its binding site sequence and context. Nucleic Acids Res. 46, 1513–1524 (2018).

49. Engler, C., Gruetzner, R., Kandzia, R. & Marillonnet, S. Golden gate shuffling: a one-pot DNA shuffling method based on type IIs restriction enzymes. PLoS One 4, e5553 (2009).

50. Thodey, K., Galanie, S. & Smolke, C. D. A microbial biomanufacturing platform for natural and semisynthetic opioids. Nat. Chem. Biol. 10, 837–44 (2014).

51. Gibson, D. G. et al. Enzymatic assembly of DNA molecules up to several hundred kilobases. Nat. Methods 6, 343–5 (2009).

52. Ryan, O. W. et al. Selection of chromosomal DNA libraries using a multiplex CRISPR system. Elife 3, (2014).

53. Kotopka, B. J. & Smolke, C. D. Production of the cyanogenic glycoside dhurrin in yeast. Metab. Eng. Commun. 9, e00092 (2019).

54. Engel, S. R. et al. The reference genome sequence of Saccharomyces cerevisiae: then and now. G3 (Bethesda). 4, 389–98 (2014).

55. Zhang, J., Kobert, K., Flouri, T. & Stamatakis, A. PEAR: a fast and accurate Illumina Paired-End reAd mergeR. Bioinformatics 30, 614–20 (2014).

56. Naughton, B. nw_align: Needleman-Wunsch basic implementation. (2019).

57. Kingma, D. P. & Ba, J. Adam: A Method for Stochastic Optimization. (2014).

58. Untergasser, A. et al. Primer3--new capabilities and interfaces. Nucleic Acids Res. 40, e115 (2012).

